# Discovery of target genes and pathways of blood trait loci using pooled CRISPR screens and single cell RNA sequencing

**DOI:** 10.1101/2021.04.07.438882

**Authors:** John A. Morris, Zharko Daniloski, Júlia Domingo, Timothy Barry, Marcello Ziosi, Dafni A. Glinos, Stephanie Hao, Eleni P. Mimitou, Peter Smibert, Kathryn Roeder, Eugene Katsevich, Tuuli Lappalainen, Neville E. Sanjana

## Abstract

The majority of variants associated with complex traits and common diseases identified by genome-wide association studies (GWAS) map to noncoding regions of the genome with unknown regulatory effects in *cis* and *trans*. By leveraging biobank-scale GWAS data, massively parallel CRISPR screens and single cell transcriptome sequencing, we discovered target genes of noncoding variants for blood trait loci. The closest gene was often the target gene, but this was not always the case. We also identified *trans*-effects networks of noncoding variants when *cis* target genes encoded transcription factors, such as *GFI1B* and *NFE2*. We observed that GFI1B *trans*-target genes were enriched for GFI1B binding sites and fine-mapped GWAS variants, and expressed in human bone marrow progenitor cells, suggesting that GFI1B acts as a master regulator of blood traits. This platform will enable massively parallel assays to catalog the target genes of human noncoding variants in both *cis* and *trans*.

## Introduction

A major goal for the study of common diseases is to identify causal genes, which can inform biological mechanisms and drug targets for these diseases. To this end, genome-wide association studies (GWAS) have identified thousands of single nucleotide polymorphisms (SNPs) or small nucleotide insertions or deletions associated with disease outcomes or disease-relevant phenotypes. However, since these associations are nearly always found in noncoding regions, identifying their target genes remains elusive. Many recent studies have used statistical fine-mapping to first identify plausibly causal GWAS variants and then combined fine-mapped variants with functional genomics data to identify candidate *cis*-regulatory elements (cCREs) and their putative target genes *(1–6)*.

However, a key missing element is the ability to move beyond association and test the impact of perturbations at cCREs on putative target gene expression at scale. Recent studies have performed CRISPR-based mutagenesis, silencing or activation screens of noncoding regulatory elements to identify target genes *(7–12)*, but no study to date has harnessed high-throughput genome engineering to study multiple GWAS variants in single cells. Here, we present Systematic Targeting and Inhibition of Noncoding GWAS loci with single-cell sequencing (STING-seq), an approach to identify GWAS target genes. STING-seq first prioritizes cCREs by functional annotation and overlap with fine-mapped GWAS variants and then tests for gene regulatory function using pooled CRISPR inhibition (CRISPRi) and single-cell RNA-sequencing (**Fig. 1A**). We demonstrate the utility and high yield of this approach by identifying target genes in *cis* and/or *trans* for 37 fine-mapped noncoding variants for 27 loci, out of 88 tested variants for 56 loci, associated across 29 blood trait GWASs from UK Biobank.

**Figure 1.**
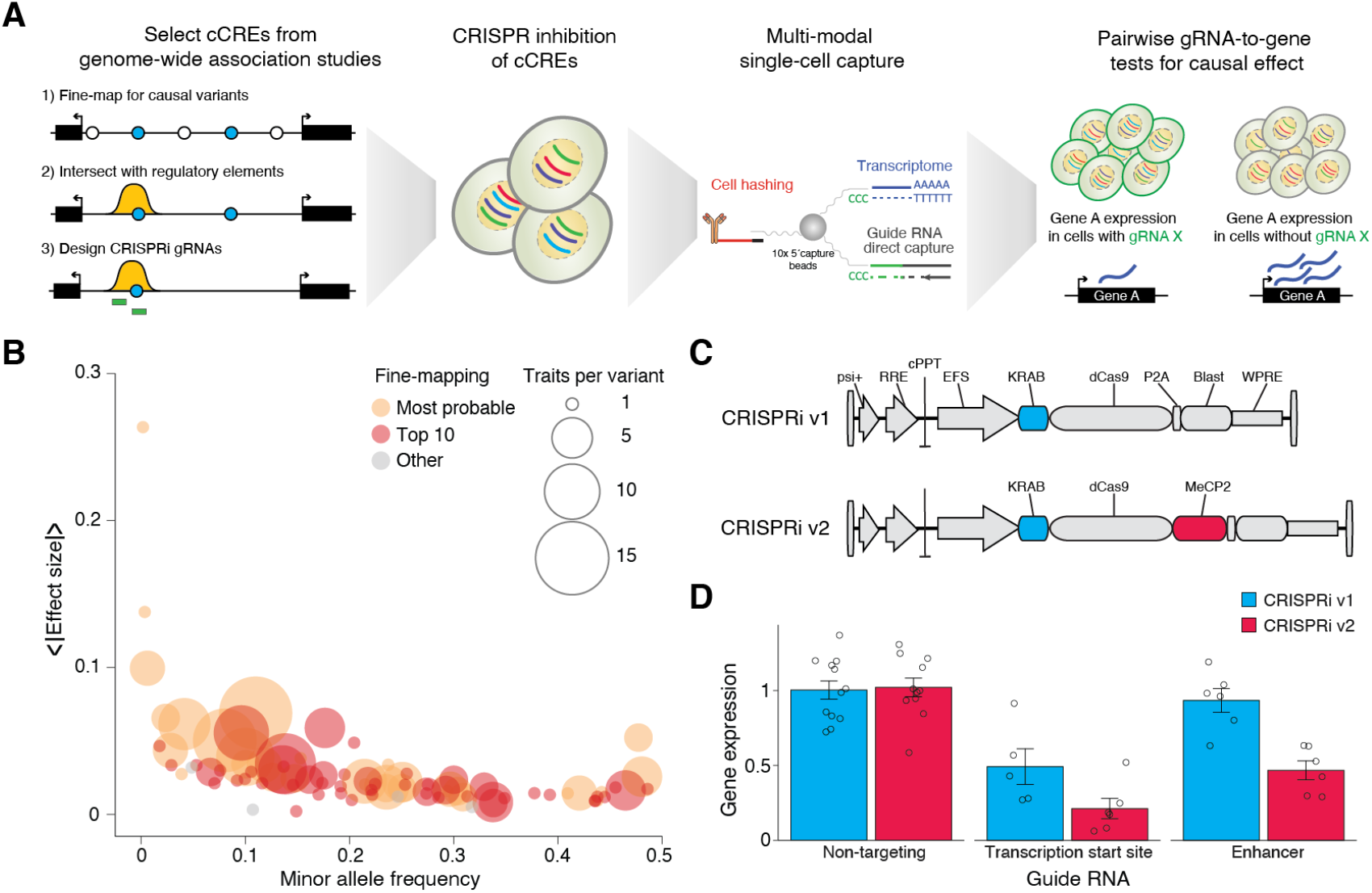
Overview of Systematic Targeting and Inhibition of Noncoding GWAS loci with single-cell sequencing (STING-seq). (**A**) STING-seq pipeline for perturbation and single-cell analysis of human genetic variants from genome-wide association studies (GWAS). Briefly, plausibly causal single-nucleotide polymorphisms (SNPs) are identified via statistical fine-mapping of GWAS. After further refinement of candidate *cis*-regulatory elements (cCREs) using key molecular hallmarks of regulatory elements, CRISPR guide RNAs (gRNAs) are designed to target cCREs and lentivirally transduced at a high multiplicity-of-infection into human cells. Using multi-modal single-cell sequencing, target genes for GWAS SNPs are identified using pairwise gRNA-to-gene tests. (**B**) Distribution of minor allele frequency (MAF) and the average absolute effect size for 88 targeted GWAS SNPs across 29 blood traits in UK Biobank (*n* = 361,194 participants). The absolute effect size is plotted for each SNP, averaged across all fine-mapped phenotypes, and the variant’s ranking among targetable variants is given by the color. (**C**) Lentiviral CRISPR inhibition (CRISPRi) with a single effector domain (KRAB, v1) or dual effector domains (KRAB and MeCP2, v2). (**D**) Digital PCR gene expression in human erythrocyte cells (K562) by targeting the transcription start sites (TSS) and known enhancers of three genes (*MRPS23, SLC25A27* and *FSCN1)* with either CRISPRi v1 or v2. We observed a stronger inhibition when targeting enhancers with the CRISPRi v2 system and used this system for STING-seq. Error bars are s.e.m.

## Results

### STING-seq library design using fine-mapped GWAS

We elected to study blood cell traits due to their high polygenicity and links to multiple common diseases *(13–15)*, and the large number of genotyped individuals available in biobank-scale data repositories with measured blood traits. For our study, we examined 29 blood trait GWASs in the UK Biobank, including traits from platelets, red blood cells, white blood cells and reticulocytes (**Table S1A**). For statistical fine-mapping, we first processed summary statistics from the 29 blood trait GWASs with GCTA-COJO *(16, 17)* to define 1 megabase (Mb) regions surrounding conditionally independent lead variants at *P*_*j*_ < 6.6×10^−9^, a suitable significance threshold for biobank-scale GWAS data *(18)* (**Table S1B**). For each 1 Mb region, we identified plausibly causal variants using FINEMAP, a shotgun stochastic search algorithm *(19)*. We identified a median of 469 conditionally independent lead variants and 3,335 fine-mapped variants (93.2% non-coding) per GWAS (**Table S1A**). For our STING-seq study, we selected a group of 88 target variants in 56 loci (1 Mb each) from different GWASs by intersecting fine-mapped noncoding variants with biochemical hallmarks of enhancer activity, such as chromatin accessibility (ATAC-seq and DNAse hypersensitivity) and canonical histone modifications (H3K27ac ChIP-seq) from human erythroid progenitor cells (K562 cells). Additionally, we required each selected variant to be distal (over 1 kb) from the transcription start site of any gene expressed in K562 cells (**Table S1C**). The variants covered a broad range of minor-allele frequencies and effect sizes and were often the highest probability variant in a fine-mapped GWAS locus (32 variants) or among the 10 most probable variants (52 variants) (**Fig. 1B**).

Before designing the CRISPR library, we sought to optimize the CRISPRi platform for our study. To date, all pooled CRISPRi screens have used a nuclease-null Cas9 (dCas9) fused to a single transcriptional repressor (KRAB) to silence specific genome elements. Recently, a dual-repressor MeCP2-dCas9-KRAB system was shown to increase repression using a transient transfection approach *(20)*. We cloned both CRISPRi repressors into a uniform lentiviral vector (Fig. 1C), transduced K562 cells and selected for transduced cells with blasticidin. We then transduced each CRISPRi cell line with guide RNAs (gRNAs) targeting the transcription start sites (TSS) of three genes (*MRPS23, CTSB* and *FSCN1)* and selected for transduced cells via puromycin selection. Using digital PCR, we found that MeCP2-dCas9-KRAB resulted in 57% greater repression of gene expression than dCas9-KRAB (**Fig. 1D, Fig. S1A-C, Table S2**). We also targeted previously described CREs for each of the 3 genes and found consistently greater repression with MeCP2-dCas9-KRAB (49% greater repression) *(7)*. Based on the increased activity at both proximal and distal regulatory elements, we proceeded with the dual-effector CRISPRi platform (“CRISPRi v2”).

To study the 88 selected variants and their cCREs, we designed a library with 176 gRNAs designed to target each variant with 2 gRNAs. We computationally optimized the 176 gRNAs to minimize off-target activity using established gRNA design tools *(21)* (**Table S3A**). We also included three sets of controls: 12 non-targeting (negative) control gRNAs from the GeCKO v2 library *(22)*, 12 positive control gRNAs to target the TSS of six genes that are highly-expressed in K562 cells, *CD46, CD52, HSPA8, NMU, PPIA* and *RPL22 (23)*, and 10 gRNAs to target the TSS of *CD55* that were used as an in-line control to estimate the average number of gRNAs per cell (multiplicity of infection, MOI) (**Table S3A**).

### STING-seq identifies *cis-*effects upon cCRE perturbation

We transduced K562 cells with the pooled library virus at a high MOI (multiple CRISPRi perturbations per cell) and verified the high MOI transduction via flow cytometry for CD55 after 10 days (**Fig. S2A-C**). We then performed Expanded CRISPR-compatible Cellular Indexing of Transcriptomes and Epitopes with sequencing (ECCITE-seq) to simultaneously capture CRISPRi gRNAs, transcriptomes and antibody-derived tags from single cells *(24, 25)*. The antibody-derived tags enabled us to recover greater numbers of single cells through droplet superloading and doublet detection.

After sequencing the ECCITE-seq library, we isolated cells with high-quality transcriptomes by eliminating cells with low total transcriptome reads and those with excessive mitochondrial read contamination. We also defined a minimum threshold of unique molecular identifiers (UMIs) for gRNA assignment (5 UMIs) beyond which there was minimal change in the number of gRNAs or cells detected (**Fig. S3A, B**). After these filters, we recovered 9,343 single cells with a median of 9 gRNAs per cell (**Fig. 2A, Table S3B**). Each gRNA was found in a median of 342 cells (**Fig. 2B**). To perform gRNA-to-gene-expression pairwise testing, we recently developed a conditional resampling approach (SCEPTRE) that yields state-of-the-art calibration on CRISPR single-cell datasets *(26)*. We applied SCEPTRE to all targeting gRNAs, performing 1,261 pairwise tests with a median of 7 genes per variant-targeting gRNA located within 500 kb of each tested SNP.

**Figure 2.**
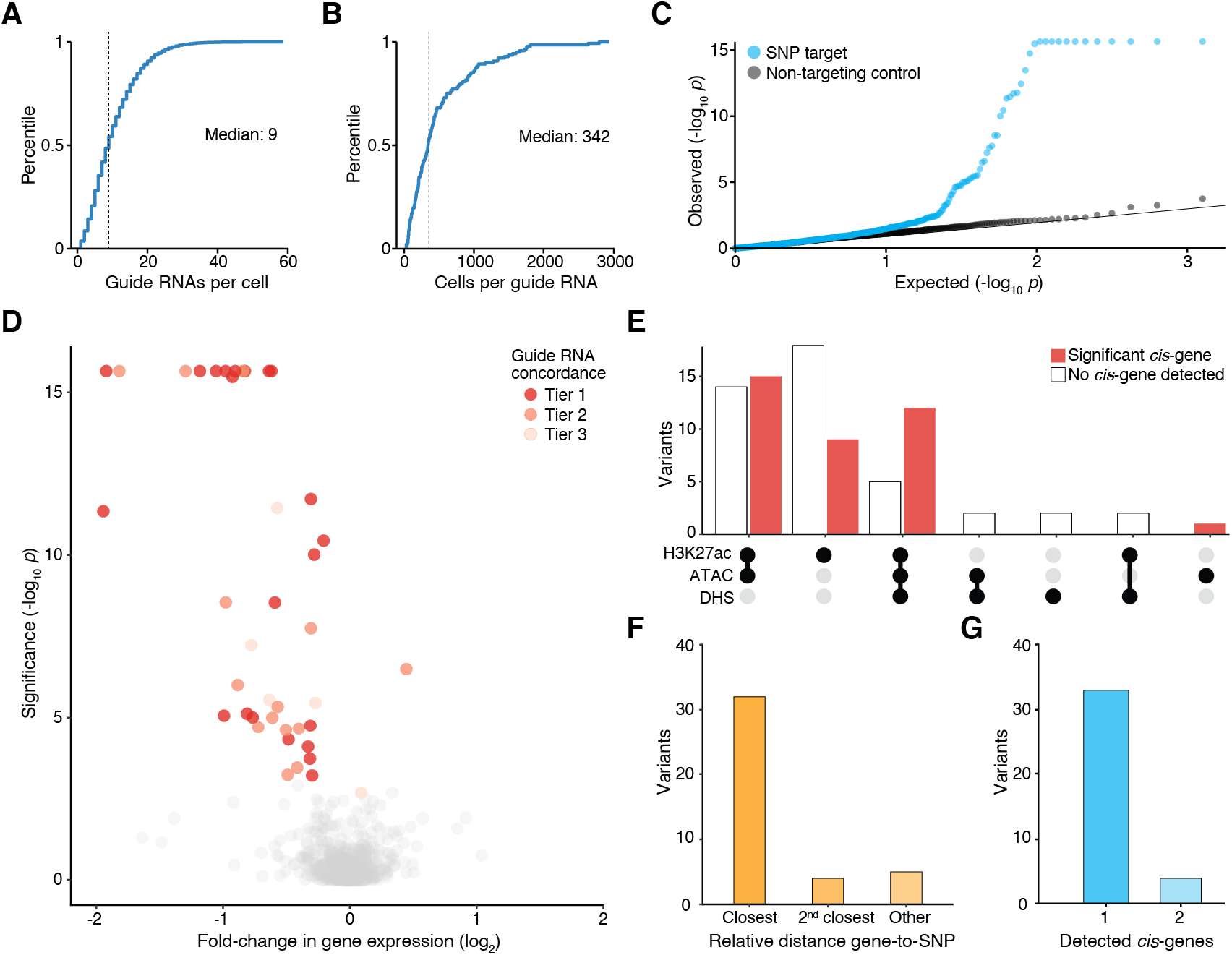
Significant *cis*-regulatory effects across 88 targeted blood trait GWAS variants. (**A**) Cumulative frequency distribution (CFD) of integrated gRNAs per K562 cell. (**B**) CFD of the number of K562 cells with each integrated gRNA. (**C**) Quantile-quantile plot of 1,261 gRNA-to-gene pairs tested in *cis* (8.9 ± 6.1 genes tested within 500 kb per gRNA). 56 gRNA-to-gene pairs tested were significant (5% FDR). Non-targeting gRNA-to-gene pairs were randomly sampled from the same set of genes tested in *cis* for targeting gRNAs. (**D**) Volcano plot of the gRNA-to-gene pairs tested in *cis*, showing the most significant gRNA per gene by significance tiers. 20 genes were identified for Tier 1 (5% FDR for the most significant gRNA and nominal *p* < 0.05 for the other gRNA), 12 genes were identified for Tier 2 (5% FDR for the top gRNA and concordant effect direction for the other gRNA) and 5 genes were identified at Tier 3 (5% FDR for the top gRNA). In total, 33 target genes were identified for 37 variants. (**E**) Targeted variants with and without a significant *cis*-gene detected. Shown below are whether the indicated variants overlap different functional hallmarks of enhancer activity in K562 cells. (**F**) Rank of distance to gene for significant variant-gene pairs. (**G**) Number of significant *cis*-genes detected per variant.

We observed good calibration for our results when compared to testing for differential expression between non-targeting gRNAs and the same 586 genes tested for *cis*-regulatory effects (**Fig. 2C, Table S3C**). None of the non-targeting gRNA-gene pairs were significant (5% false discovery rate, FDR). In contrast, at the same 5% FDR threshold, positive control gRNAs targeting the TSSs of six genes were all associated with significant decreases in expression (**Fig. S3C, Table S3D**).

We used the same threshold to identify STING-seq CREs with significant *cis*-regulatory effects and found that 37 of the variants met this criteria.

Since we designed our STING-seq library with two independent CRISPRi gRNAs targeting each variant, we also quantified the concordance between genetic perturbations by assigning variants to one of three Tiers (**Fig. 2D, Table S3E**). In all Tiers, we required that at least one gRNA targeting the CRE to have a significant change in gene expression in *cis* (5% FDR). The majority of CREs with significant *cis*-regulatory effects (21 variants) had their second gRNA reach nominal significance with a concordant change in gene expression (SCEPTRE *p* < 0.05) (Tier 1). For Tier 2, we had 12 variants where the second gRNA had a concordant change in gene expression with the first gRNA but was not nominally significant. And, in Tier 3, we identified four variants where the second gRNA was captured in fewer than 30 cells and hence not tested. In total, we identified 33 target genes for 37 variants from blood trait GWASs. This includes loci where multiple CREs modulate the same gene or where a single CRE modulates more than one gene in *cis*.

Targeted variants that overlapped with H3K27ac ChIP, ATAC and DHS from K562 were enriched for significant *cis*-genes, compared to variants outside these annotations (odds ratio = 3.36, Fisher’s exact test *p* = 0.03) (**Fig. 2E**). For the majority of significant *cis*-genes, they were also the closest gene (distance to the TSS); however, there were four significant *cis*-genes that were the second closest, and five that were further away (**Fig. 2F**). For 33 variants, we identified a single significant gene in *cis* and, for four variants, we found two significant genes in *cis* (**Fig. 2G**). Of the 33 variants with eQTL data (eQTL Catalogue), we found that six variants had the same *cis*-eQTL target genes as their STING-seq target genes. This shows that STING-seq uncovers *cis*-target genes not captured by eQTL mapping *(27)* (**Table S3F**).

Among the loci with significant target genes in *cis*, rs4526602 was the top ranked and most significantly associated SNP to the monocyte percentage blood trait at its locus (**Fig. 3A**). The SNP is in an intergenic region, between the gene bodies of *AIM1L* and *CD52*, and the gene with the closest TSS to the SNP is *UBXN11*. Given this position, it is unclear which of these genes (if any) might be the target gene of the SNP locus. When testing for changes in gene-expression in *cis* using the most abundant gRNA (1,278 cells), we found that *CD52* was the only significantly decreased gene (**Fig. 3B**, log_2_ fold-change = -1.92, SCEPTRE *p*-value = 4.5×10^−12^).

**Figure 3.**
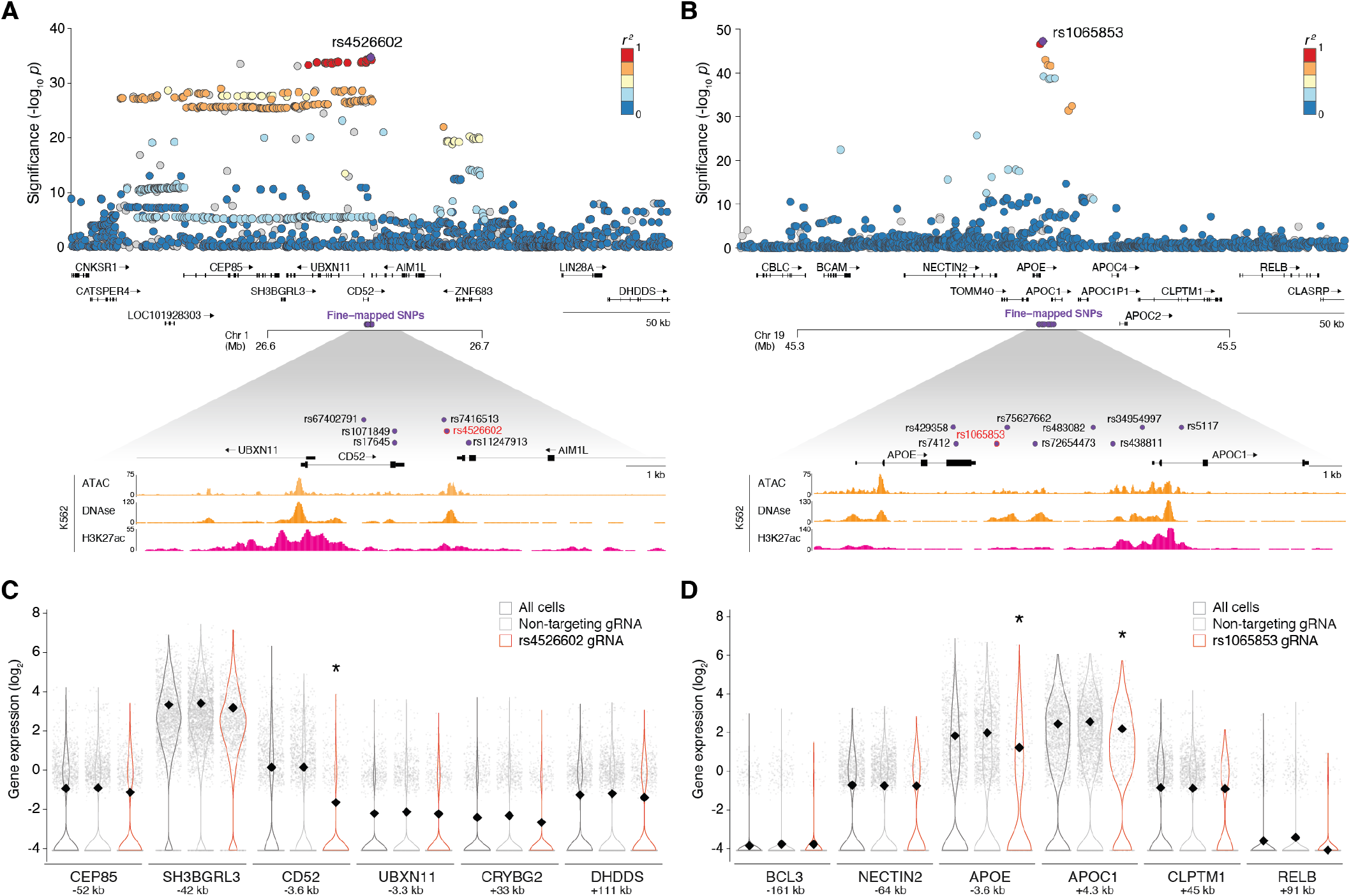
Perturbation of GWAS SNPs with *cis*-regulatory effects at the *CD52* and *APOE/ APOC1* loci. (**A**) LocusZoom plot for monocyte percentage GWAS. The lead SNP that fine-mapped as plausibly causal (rs4526602) is indicated in red. (**B**) LocusZoom plot for percentage of immature red blood cells GWAS. The lead SNP that fine-mapped as plausibly causal (rs1065853) is indicated red. (**C**) Normalized single-cell gene expression for the top rs4526602-targeting gRNA (*n* = 1,278 cells), for non-targeting gRNAs, and all cells. All genes with detected expression are shown. *CD52* is the only gene with a significant (* 5% FDR) change in expression in *cis*. (**D**) Normalized single-cell gene expression for the top rs1065853-targeting gRNA (*n* = 405 cells), for non-targeting gRNAs, and all cells. All genes with detected expression are shown. *APOE* and *APOC1* are the only genes with a significant change in expression in *cis*.

Among fine-mapped variants with two *cis*-genes detected via STING-seq, rs1065853 was the top ranked and most significantly associated SNP to an immature red blood cell percentage trait (high light scatter reticulocyte percentage) at its locus (**Fig. 3C**). This SNP maps to an intergenic region, between the gene bodies of *APOE* and *APOC1*, with *APOE* being the closest gene and also has significant associations with high and low density lipoprotein levels *(28)*. Upon testing against genes in *cis* with the top gRNA (captured in 405 cells), both *APOE* (log_2_ fold-change = -0.63, SCEPTRE *p*-value = 2.8×10^−6^) and *APOC1* (log_2_ fold-change = -0.27, SCEPTRE *p*-value = 3.5×10^−6^) had significantly decreased expression (**Fig. 3D**). Previous studies have shown that *APOE* and *APOC1*, which encode apolipoproteins E and C1, influence blood lipids and diverse ailments including cardiovascular disease and Alzheimer’s disease *(29, 30)*. Since these genes work in a coordinated fashion to regulate lipid metabolism *(31)*, the co-regulation of these genes by the rs1065853 locus is an especially interesting observation of regulatory pleiotropy that may contribute to trait associations.

We also examined loci with several fine-mapped variants near a single gene of interest. At the *PTPRC* locus, we targeted nine variants that were significant across 10 GWASs (**Fig. S4A, Table S1D**) and mostly not in strong LD as quantified by pairwise R^2^ from 1000 Genomes *(32)* (**Fig. S4B**). The nine variants mapped to distinct cCREs, one was upstream of the *PTPRC* TSS and the remaining eight were in the first intron (**Fig. S4C**). We observed the strongest repression of *PTPRC* when targeting the two SNP-identified CREs closest to the TSS (**Fig. S4D**), and significant effects for four other SNP-CREs. Of the three nonsignificant targeted SNPs, two are in high LD (R^2^ ≥ 0.95) for significant targeted SNPs, suggesting that these may be nonfunctional LD proxies. For all of these variants, *PTPRC* was the only significant target gene and thus very likely the causal GWAS gene (**Table S3E**). These results suggest high allelic heterogeneity driven by multiple independent regulatory variants in distinct CREs affecting *PTPRC* expression. Interestingly, these variants have highly distinct blood trait associations (**Fig. S4A**), indicating that while regulatory effects from distinct CREs can converge at the level of the expression of a single gene, their downstream phenotype effects can vary.

### Inhibition of a CRE that modulates *GFI1B* expression leads to widespread *trans*-effects

To understand the impact of GWAS CREs on global gene expression, we tested all gRNAs for *trans*-effects by performing transcriptome-wide differential gene expression tests against all 10,324 genes expressed in at least 5% of cells. We applied a strict 1% false discovery rate to identify significant genes in *trans* (**Fig. 4A**) and again saw good calibration with non-targeting gRNAs (**Table S3C**). We observed a striking number of significant *trans*-effects for gRNAs targeting three different SNPs: rs524137, rs73660574 and rs79755767 (**Fig. 4A, Fig. S5, Table S3G-I**). We found that rs524137 and rs73660574 were in significant CREs for *GFI1B* (**Fig. 4B**) and that rs79755767 was in a significant CRE for *NFE2* (**Fig. S6A**). GFI1B and NFE2 are both transcription factors that play a key role in the differentiation of hematopoietic stem cells *(33)*.

**Figure 4.**
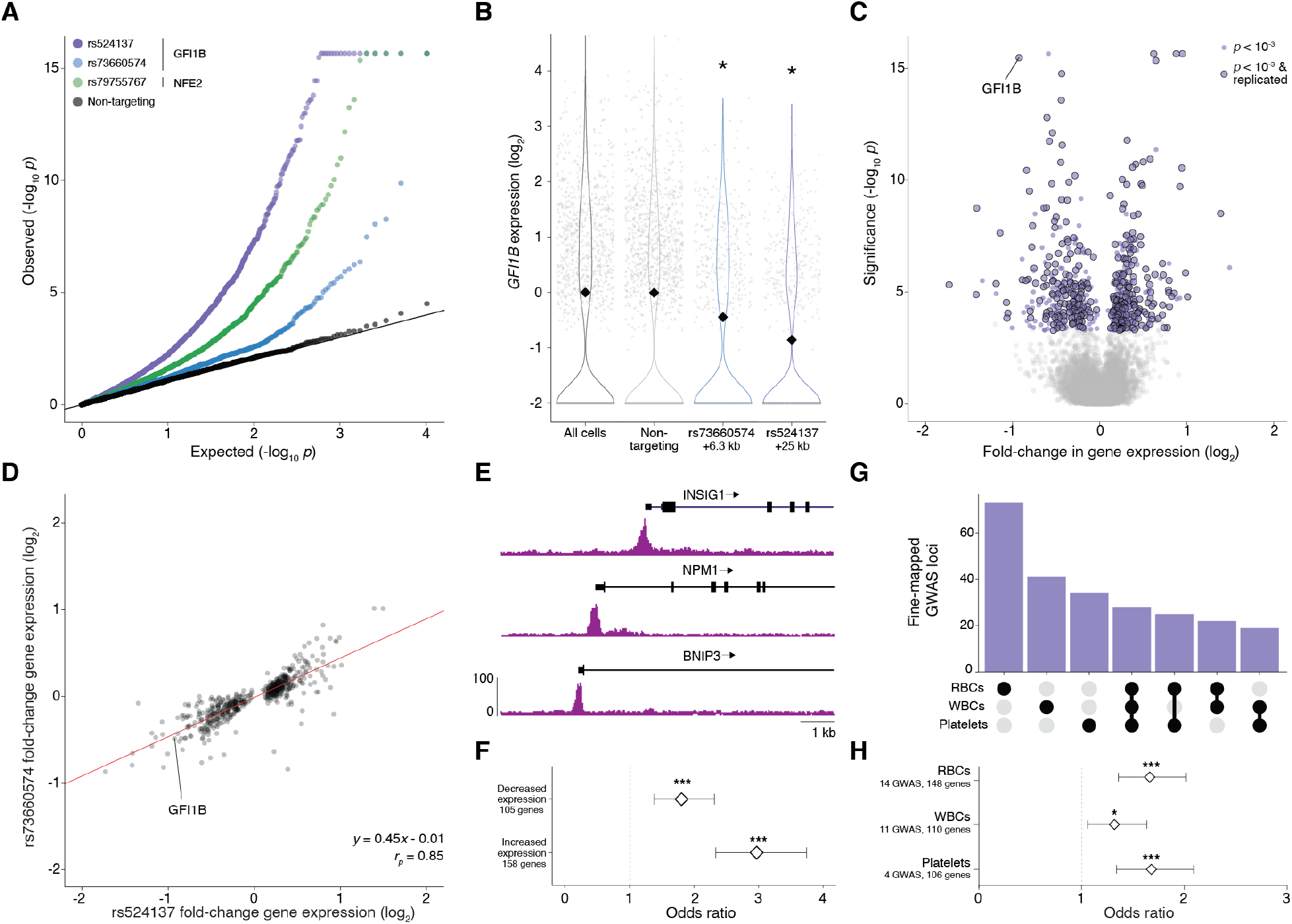
Perturbation of *cis*-regulatory elements of the transcription factor GFI1B reveal a *trans*-regulatory network of genes that impact diverse blood cell traits. (**A**) Quantile-quantile plot of transcriptome-wide gRNA-to-gene pairs tested in *trans* (10,319 genes tested per gRNA). Shown are the top gRNAs for rs524137 (intergenic CRE for *GFI1B*), rs73660574 (intronic CRE for *GFI1B*) and rs79755767 (intergenic CRE for *NFE2)* and non- targeting gRNA-to-gene pairs randomly sampled from transcriptome-wide tests. (**B**) *GFI1B* expression was significantly decreased (* 5% FDR) when targeting either of two *cis* SNPs, rs524137 and rs79755767. (**C**) Volcano plot of top rs524137-targeting gRNA-to-gene pairs in *trans*. We identified 569 unique genes for the rs524137-*GFI1B trans*-regulatory network. (**D**) Scatter plot of expression fold-change for the 569 rs524137-*GFI1B trans*-regulatory genes when perturbing rs524137 and rs79755767. There is a high correlation between these perturbations (*r*_*p*_ = 0.85) and the dosage effect (slope = 0.45) is in agreement with the difference in rs524137 and rs79755767 effects on *GFI1B* expression (Δlog_2_ fold-change = 0.44). (**E**) GFI1B ChIP-seq at the transcription start sites of three rs524137-*GFI1B trans*-regulated genes (*INSIG1, NPM1* and *BNIP3)*. (**F**) rs524137-*GFI1B trans*-regulated genes were significantly enriched for *GFI1B* ChIP-seq peaks. Genes with increased expression [odds ratio (OR) = 2.96, Fisher’s exact test *p*-value (*p*) = 5.5×10^−20^] were more strongly enriched than genes with decreased expression (OR = 1.79, *p* = 9.3×10^−6^). (**G**) Overlap of genes closest to fine-mapped GWAS variants, grouped by the cell type of the GWAS trait (platelets, white blood cells or red blood cells), for genes that are also rs524137-*GFI1B trans*-regulated genes. (**H**) rs524137-*GFI1B trans*-regulated genes were enriched for 29 blood trait GWAS loci, grouped together as platelet genes (OR = 1.68, *p* = 8.5×10-6), red blood cell genes (RBC; OR = 1.66, *p* = 6.0×10^−7^) and white blood cell genes (WBC; OR = 1.32, *p* = 0.01). In (F) and (H), * indicates *p* < 0.05, ** indicates *p* < 0.01 and *** indicates *p* < 0.001.

For rs524137, an intergenic enhancer of *GFI1B* that is located 11.5 kb 3’ of the gene, we observed 569 differentially expressed genes (**Fig. 4C**). By analyzing data from a prior CRISPRi-based study that also targeted this genomic locus and reported the *cis* effect on *GFI1B (7)*, we found that 321 of these differentially-expressed genes replicated independently (**Fig. 4C**). For the rs73660574-identified CRE, a weaker, intronic enhancer for *GFI1B*, we observed 28 differentially expressed genes; 24 of these 28 differentially expressed genes (including *GFI1B*) were also found when perturbing the rs524137 locus (**Fig. S6B**). Thus, perturbing either rs73660574-or rs524137-identified CREs led to changes in the expression of target genes of GFI1B.

To better understand the *trans*-effects of these two *GFI1B* enhancers, we examined gene-expression changes in all 569 differentially-expressed genes identified for rs524137 — and did so for both SNPs (**Fig. 4D**). For these genes, we observed a high correlation between perturbations targeting each SNP (*r* = 0.85). Even though many of the gene expression changes were more modest when perturbing rs73660574, we found a linear dosage relationship between the *cis* and *trans* regulatory effects for these two SNP loci: the dosage effect across the 569 genes (log_2_-slope = 0.45) was in agreement with the difference in rs524137 and rs79755767 effects on *GFI1B* expression (Δlog_2_ fold-change = 0.44) (**Fig. 4B**).

### *Trans*-regulated genes are enriched for transcription factor binding sites and GWAS loci

*GFI1B* encodes a transcription factor central to hematopoiesis *(33–36)*, therefore we examined GFI1B ChIP-seq data in K562 cells *(35)* near the transcription start sites of rs524137-*GFI1B trans*-regulated genes and observed GFI1B binding (**Fig. 4E, Table S4A**), with a significant enrichment across all rs524137-*GFI1B trans*-regulated genes (**Fig. 4F, Table S4B**). As expected given *GFI1B*’s known role as a transcriptional repressor *(36)*, genes with increased expression were more strongly enriched [odds ratio (OR) = 2.96, Fisher’s exact test *p*-value (*p*) = 5.5×10^−20^] than genes with decreased expression (OR = 1.79, *p* = 9.3×10^−6^). Among significant rs73660574 *trans*-regulated genes for GFI1B binding sites only the genes with increased expression were significantly enriched (OR = 5.22, *p* = 5.6×10^−3^, **Fig. S6C, Table S4B**).

The *NFE2* CRE has significant *trans*-effects (rs79755767) for 134 genes (**Fig. S6D**), supported by a previous study that reported its *trans*-effects *(8)* that are potentially explained by *NFE2*’s role as a hematopoietic transcription factor important for platelet and megakaryocyte development *(33, 37)*. Using NFE2 ChIP-seq data from K562 cells *(38)*, we observed an enrichment for rs79755767-*trans* regulated genes (OR = 6.12, *p* = 4.6×10^−8^), including both genes with increased (OR = 5.94, *p* = 1.3×10^−4^) and decreased expression (OR = 6.25, *p* = 9.5×10^−5^) (**Fig. S6E, Table S4A-B**). This was not surprising, as NFE2 is a member of a complex of transcription factors and can act as either a gene activator or repressor *(39)*. These findings suggest that perturbing CREs for TFs that are putatively affected by fine-mapped noncoding GWAS variants can reveal genes that are the primary targets for TF binding and secondary targets in a regulatory network.

Next, we hypothesized that GFI1B and NFE2 *trans*-regulatory networks discovered by STING-seq may have an overall role in blood trait GWASs, beyond the variants affecting these TFs in *cis*. We constructed a set of putatively causal genes for each of the 29 GWASs by selecting the closest genes to fine-mapped variants of GWAS loci, and then grouped them by their blood cell trait, generating gene sets for platelets, red blood cells (RBCs) and white blood cells (WBCs) (**Table S4A**). Analyzing genes that overlap rs524137-*GFI1B trans*-regulated genes, we discovered that the gene sets of these three GWAS cell-types were mostly distinct (**Fig. 4G**). The rs524137-*GFI1B trans*-regulatory genes were broadly enriched for blood cell GWAS traits, including platelet genes (OR = 1.68, p = 8.5×10^−6^), RBC genes (OR = 1.66, p = 6.0×10^−7^) and WBC genes (OR = 1.32, p = 1.3×10^−2^) (**Fig. 4H, Table S4B**). We also observed an enrichment of rs79755767-*NFE2 trans*-regulatory networks for red blood cell genes (OR = 1.76, *p* = 6.2×10^−3^) (**Fig. S6F-G, Table S4B**), indicating that the known role of *GFI1B* and *NFE2* in hematopoiesis and cell differentiation is mediated by their effects on regulatory networks. Enrichment of independent blood cell trait GWAS loci among *GFI1B* and *NFE2 trans*-regulated genes provides strong support for this.

### *Trans*-regulated genes cluster by biological role and cell type

Given the breadth of rs524137-*GFI1B trans*-regulated genes and their enrichment for GWAS loci, we next analyzed the structure of this regulatory network and its contribution to blood trait GWAS and underlying biology. Using gene co-expression and clustering analyses with our K562 single-cell data on the 569 identified rs524137-*GFI1B trans*-regulated genes, we identified three major gene clusters (A, B and C) (**Fig. 5A**). Cluster A, which represents approximately half of the genes with increased expression upon repression of the *GFI1B* CRE with STING-seq (**Fig. 5A, Table S4A**), was the most strongly enriched for *GFI1B* binding sites (OR = 4.02, *p* = 7.4×10^−8^) (**Fig. 5B, Table S4B**). Cluster B, which includes the other half of genes with increased expression, and Cluster C, which consists almost entirely genes with decreased expression, were also enriched for GFI1B binding sites, although less so (OR = 2.00, *p* = 6.2×10^−5^ and OR = 1.82, *p* = 7.0×10^−6^, respectively) (**Fig. S7A, Fig. 5C, Table S4B**). Cluster C was strongly enriched for putative GWAS genes for platelet and RBC traits (OR = 1.76, *p* = 7.1×10^−4^ and OR = 2.02, *p* = 1.3×10^−6^, respectively) (**Fig. 5C, Table S4B**), which was consistent with Gene Ontology enrichments in this cluster for heme biosynthesis (**Table S4C**). Given that Cluster A was the most strongly enriched for GFI1B binding, but Cluster C was the most strongly enriched for GWAS loci and contained known genes for blood biochemistry, we inspected these two clusters further in primary cells to inform what the separate or shared functions of these clusters may be.

**Figure 5.**
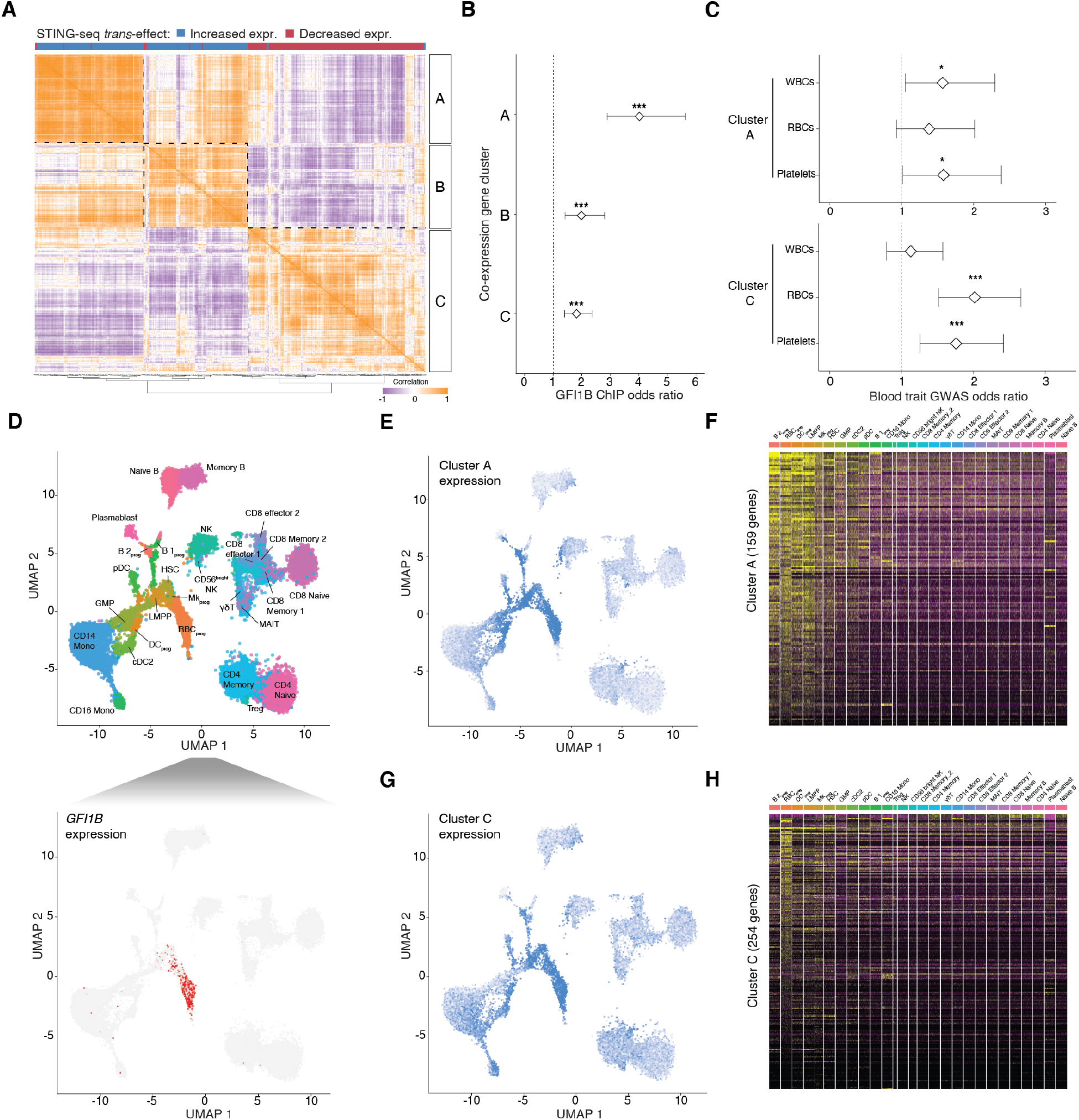
Gene co-expression analysis identifies subnetworks of GFI1B target genes that are expressed in multiple hematopoietic progenitors. (**A**) Co-expression matrix of rs524137-*GFI1B* trans-regulatory genes in K562 cells with hierarchical clustering. Three clusters (A, B and C) are indicated and bar underneath indicates if the genes had increased (*blue*) or decreased (*red*) expression upon perturbing the *GFI1B* CRE. (**B**) Clusters A, B and C all have significant enrichment for GFI1B binding sites, with Cluster A having the strongest enrichment. (**C**) Clusters A and C enrichments for genes mapping closest to fine-mapped variants for WBC, RBC and platelet GWAS traits. (**D**) UMAP plot of human bone marrow cells from 35 Human Cell Atlas donors aligned into a common coordinate space. Labels and colors indicate distinct cell types (B 2_prog_: progenitor B-2 cells; RBC_prog_: red blood cell progenitors; DC_prog_: dendritic cell progenitors; full list in **Table S4D**). Below *GFI1B* single cell expression projected onto the UMAP plot is shown, indicating that *GFI1B* is most highly expressed in red blood cell progenitors and megakaryocyte progenitors. (**E**) Mean expression of Cluster A genes (*n* = 159 genes) per single cell, projected onto the UMAP plot in (*C*). The cell types with the highest Cluster A expression patterns are bone marrow progenitor cells, whereas expression in differentiated cells is lower. (**F**) Expression of genes in Cluster A by cell type in 35 Human Cell Atlas bone marrow donors. (**G**) Mean expression of Cluster C genes (*n* = 254 genes) per single cell, projected onto the UMAP plot in (*C*). The cell types with the highest Cluster C expression patterns are RBC progenitors and megakaryocyte progenitors. (**H**) Expression of genes in Cluster C by cell type in 35 Human Cell Atlas bone marrow donors. * indicates *p* < 0.05, ** indicates *p* < 0.01 and *** indicates *p* < 0.001.

We integrated the *GFI1B* co-expression network with Human Cell Atlas single-cell RNA-sequencing from 35 bone marrow donors *(40, 41)*, as bone marrow includes a rich sample of multipotent progenitor cells crucial for hematopoiesis (**Fig. 5D**). First, we confirmed the expression of *GFI1B* was limited to progenitor cells for RBCs and megakaryocytes, which produce platelets *(33)* (**Fig. 5D**). Genes from Cluster A were expressed in virtually all bone marrow progenitor cells, but not in differentiated cells (**Fig. 5E-F, Table S4D**). Together with Cluster A’s high enrichment of GFI1B binding sites and increased expression upon inhibiting *GFI1B* (a repressor), this suggests that these genes are the primary targets of *GFI1B* to suppress differentiation in non-erythrocytic and non-megakaryocytic pathways *(36)*. Cluster B genes were highly expressed in granulocyte-monocyte progenitors (GMP) and their differentiated cell types, including monocytes and dendritic cells (**Fig. S7B-C, Table S4D**). For Cluster C, we saw expression in RBC progenitors and megakaryocyte progenitors (**Fig. 5G-H**). We summarized the findings for the three clusters (**Table S4E**) and computed their mean expression per cell across all bone marrow cell types (**Fig. S7D**). These results suggest that Cluster C genes, which had decreased expression upon inhibiting *GFI1B* via STING-seq and are enriched for genes in RBC and platelet GWAS loci as well as in the heme biosynthesis pathway, are secondary targets of *GFI1B* to differentiate progenitor cells down the RBC and megakaryocytic lineages. That we identified these *trans*-regulatory networks in a homogenous blood progenitor-like cell type (K562) demonstrates the utility of STING-seq in studying diverse effects of CREs on target genes. GWAS-identified CREs can impact entire regulatory networks in *trans*, providing insight into molecular mechanisms of genetic effects on complex traits.

### Estimation of single cells needed to apply STING-seq to other diseases and traits

Given the large number of GWASs performed over the past 15 years, with numbers of trait-associated loci per GWAS ranging from tens to thousands *(28)*, we wanted to understand the scale of cells needed to perform STING-seq under various settings. By performing statistical down-sampling experiments on the *trans*-regulatory effects identified with STING-seq, we computed the number of cells required for 80% power with high, medium and low target gene expression, and strong, medium and weak CRISPRi effects (*i*.*e*. observable log_2_ fold-change) (**Fig. S8A**). For example, for genes in the middle tertile of expression with strong CRISPRi effects, 80% power was retained with as few as 350 cells per gRNA. To extrapolate the number of single cells required for other GWASs, we took the number of independent loci for recent studies on BMI-adjusted waist-hip ratio (1077 loci) *(42)*, type 2 diabetes (568 loci) *(43)*, schizophrenia (222 loci) *(44)*, rheumatoid arthritis (101 loci) *(45)* and stroke (32 loci) *(46)*. If SNP prioritization allowed targeting of a single SNP per locus, the same design as in this study would allow search for *trans*-effects at 80% power across several hundreds of GWAS loci with approximately 50,000 single cells – which is a feasible experimental scale (**Fig. S8B**). If this analysis was performed under typical pooled CRISPR screen parameters with MOI ∼ 1, at least 500,000 cells would be necessary, highlighting the utility of multiple CRISPRi perturbations per cell in STING-seq.

## Discussion

In summary, we have developed an approach to understand noncoding human genetic variants that integrates fine-mapped GWAS data, pooled CRISPR screens and single-cell RNA-sequencing and can identify target genes for variants in both *cis* and *trans*. We demonstrated the utility of STING-seq in identifying *cis*-target genes of CREs with likely causal GWAS variants, and also describe more complex regulatory architectures with CREs regulating more than one gene in *cis* and multiple CREs targeting the same gene. Additionally, we identified variants mapping to CREs for the transcription factors *GFI1B* and *NFE2* and, through their perturbation, identify *trans*-regulatory networks and their subnetworks with distinct biological functions. The enrichment of genes in independent blood cell trait GWAS loci in these networks implies a polygenic contribution in these cellular functions that underlie diverse blood cell traits.

The first step of STING-seq — the selection of putatively causal variants GWAS variants to target — relies on prioritization approaches *(2, 6, 47–52)*. We used a straightforward annotation strategy that looked for fine-mapped variants in cCREs, and identified significant target genes for 42% of them. We considered this a high yield over previous studies that studied regulatory effects of noncoding genomic loci *(7–9, 53–60)*. The loci without detected effects can be due to the variant being a non-functional LD proxy of the real causal variant, lack of effect on the studied cell type, or simple lack of detection power. Improved fine-mapping and variant prioritization methods have the potential to improve target gene identification with STING-seq further especially in loci with high LD.

STING-seq uses a powerful and scalable CRISPRi-based approach to suppress GWAS-implicated CREs. However, functional effects of a variant cannot be directly inferred from silencing of a regulatory element, but fine-mapping of GWAS-associated variants overlapping experimentally validated CRE provide strong joint evidence of these variants being functional. Going forward, STING-seq can be combined with catalytically active Cas9, CRISPR-activation or scalable base-editing approaches *(61)*. A key feature of recent CRISPRi screens of CREs *(7, 8)*, including STING-seq, is introduction of multiple perturbations per cell. This substantially increases the number of loci that can be feasibly analyzed. While this is feasible for immortalized cell lines like K562, expanding multiple perturbations (via either high MOI transduction or innovative vector designs) to other cell lines and primary cells will be instrumental for the next stage of target gene identification and characterization for diverse GWAS traits.

Finally, our results demonstrate the power of single-cell sequencing for sensitive and scalable readout of regulatory effects of GWAS loci in *cis* and *trans*. While we have a high yield in *cis* target gene discovery, we note that identification of a *cis* gene alone with STING-seq does not prove its mechanistic causal role driving the GWAS association, nor exclude other potential causal variants, CREs and genes, including in other cell types. In loci where *cis*-effects are coupled with *trans*-network effects, STING-seq can be highly informative of potential cellular mechanisms, which also provides strong support for the causal role of the *cis*-target gene.

Altogether, the workflow we have outlined with STING-seq demonstrates a clear roadmap to identify target genes for GWAS loci in a high-throughput fashion, enabling future studies of human noncoding genome function.

## Acknowledgments

We thank the entire Sanjana and Lappalainen laboratories for support and advice. We also thank the New York Genome Center for flow cytometry resources, the NYU Biology Genomics Core for sequencing resources, E. Hoelzli for support with digital PCR, T. Stuart and H. Wessels for advice on single cell sequencing quality control and S. Kasela for help with library design.

## Author contributions

J.A.M and N.E.S. conceived the project. J.A.M., D.A.G., and N.E.S. performed the GWAS statistical fine-mapping, functional annotation and STING-seq library design. Z.D. and M.Z. developed the lentiviral constructs and monoclonal K562 cells. J.A.M. and Z.D. designed and performed lentiviral transductions. J.A.M, S.H. and E.P.M. performed ECCITE-seq. J.A.M., T.B. and E.K. performed differential gene expression testing. J.A.M. and J.D. performed co-expression analyses and human bone marrow single-cell data analyses. P.S., K.R., E.K., T.L. and N.E.S. supervised the work. J.A.M., T.L. and N.E.S. wrote the manuscript with input from all authors.

## Funding

J.A.M is supported by a Canadian Institutes of Health Research Banting Postdoctoral Fellowship. Z.D is supported by an American Heart Association postdoctoral fellowship (20POST35220040). T.L. is supported by NIH/NIMH (R01MH106842), NIH/NHGRI (UM1HG008901) and NIH/NIGMS (R01GM122924). N.E.S. is supported by NYU and NYGC startup funds, NIH/NHGRI (DP2HG010099, including a supplemental award for STING-seq), NIH/NCI (R01CA218668), NIH/NIGMS (R01GM138635), DARPA (D18AP00053), the Cancer Research Institute, the Sidney Kimmel Foundation and the Brain and Behavior Foundation.

## Competing interests

N.E.S. is an advisor to Vertex. T.L. is an advisor to Goldfinch Bio and GSK and, with equity, Variant Bio.

## Data and materials availability

Plasmids (lentiCRISPRi(v1)-Blast, lentiCRISPRi(v2)-Blast and lentiGuideFE-Puro) have been deposited with Addgene (plasmid nos. 170067, 170068, 170069). Raw and processed single cell sequencing files are available at the Gene Expression Omnibus (GEO accession number GSE171452).

## Materials and methods

### Genome-wide association studies of blood cell traits

We used genome-wide association study (GWAS) summary statistics for 29 blood cell traits from 361,194 white British UK Biobank participants: White blood cell (leukocyte) count, red blood cell (erythrocyte) count, hemoglobin concentration, hematocrit percentage, mean corpuscular volume, mean corpuscular hemoglobin, mean corpuscular hemoglobin concentration, red blood cell (erythrocyte) distribution width, platelet count, platelet crit, mean platelet (thrombocyte) volume, platelet distribution width, lymphocyte count, monocyte count, neutrophil count, eosinophil count, basophil count, lymphocyte percentage, monocyte percentage, neutrophil percentage, eosinophil percentage, basophil percentage, reticulocyte percentage, reticulocyte count, mean reticulocyte volume, mean sphered cell volume, immature reticulocyte fraction, high light scatter reticulocyte percentage and high light scatter reticulocyte count (**Table S1A**). Each GWAS was performed by fitting the following covariates to inverse normal transformed traits with linear regression models: Principal components 1 through 20, sex, age, age^2^, sex and age interaction and sex and age^2^ interaction. The summary statistics were generated by the Neale Lab (www.nealelab.is/uk-biobank).

### Statistical fine-mapping of blood cell traits

The 29 GWASs of blood cell traits were uniformly processed with a statistical fine-mapping pipeline. First, each GWAS was analyzed with GCTA-COJO v.1.93.1 *(16, 17)* to identify conditionally independent lead variants (*p*_*j*_ < 6.6×10^−9^) and define 1 Mb regions for statistical fine-mapping. All variants within 500 kb of a lead variant were analyzed with FINEMAP v.1.3.1 *(19)*, a Bayesian fine-mapping method that assigns each variant a Bayes factor for being plausibly causal. Both GCTA-COJO and FINEMAP require population-matched covariance matrices, therefore we generated these with PLINK v.2.0 *(62)*, QCTOOL v.2.0.2, BGENIX v.1.1.5 *(63)* and LDstore v.1.1 *(64)*, using a subset of 50,000 UK Biobank white British participants (UK Biobank accession code 47976). FINEMAP allows for a maximum number of causal configurations to test for each input set of variants, therefore we set the maximum to 10 causal configuration variants per fine-mapped region and excluded cases where FINEMAP failed to converge. We then retained noncoding variants with a high Bayes factor (log_10_ BF ≥ 2) for a set of causal variants. Fine-mapped variants that had more than one Bayes factor, due to being within 250 kb of multiple lead variants, had their highest value retained. Across all 29 GWASs, we identified 825 loci, separated by at least 500 kb, and 55,349 fine-mapped variants. The Variant Effect Predictor (VEP) tool *(65)* was used to identify 52,293 of which were noncoding.

### Functional annotation of causal noncoding SNPs

We integrated multiple functional genomics datasets for K562 cells. Specifically, we used DNase I hypersensitive sites (DHS) from ENCODE *(38)*, H3K27ac ChIP-seq peak calls from ENCODE, and ATAC-seq peak calls that we generated previously *(66)* to identify candidate *cis*-regulatory elements (cCREs). We used bedtools v.2.25.0 *(67)* and bedops v.2.4.3 *(68)* to identify variants mapping directly to cCREs. We also required variants to be further than 1 kb from any gene TSS. This annotation strategy identified 10,789 distinct variants mapping to 627 loci. We then selected 88 variants from 56 loci for targeting based on whether a variant was targetable and more plausibly causal than others for a given GWAS and locus by ranking FINEMAP log_10_ Bayes factors and manual inspection of loci. For the 88 selected variants, 32 were the most probable variant for at least one GWAS locus, and 52 were in the top-10 most probable variants. For the 56 loci, there was an average median of 10.5 (± 8.6) targetable SNPs. Elements of manual inspection included selecting variants that mapped to intergenic regions between gene TSSs or selecting multiple variants that map proximal to the same gene.

### Plasmid cloning for lentiviral CRISPRi and modified gRNA scaffold vectors

To generate the KRAB-dCas9 (lentiCRISPRi(v1)-Blast) and KRAB-dCas9-MeCP2 (lentiCRISPRi(v2)-Blast) plasmids, KRAB and dCas9 were PCR amplified from pCC_09 (Addgene 139094) *(69)* and the MeCP2 effector domain was synthesized as a gBlock (IDT). KRAB and MeCP2 were linked to dCas9 with flexible glycine-serine linkers and cloned into lentiCas9-Blast (Addgene 52962) *(22)*.

To generate the gRNA vector (lentiGuideFE-Puro), we digested pCC_09 with NheI and KpnI to isolate the U6 promoter and Cas9 guide RNA scaffold with the F+E scaffold modification *(70)*. After gel extraction (Qiagen 28706), we ligated this piece into NheI and KpnI-digested pLentiRNAGuide_001 (Addgene 138150) vector using T4 ligase (NEB M0202M) *(71)*.

### Cell culture and monoclonal cell line generation

HEK293FT cells were acquired from Thermo Fisher (R70007). HEK293FT (human) cells were maintained at 37°C with 5% CO2 in D10 media: DMEM with high glucose and stabilized L-glutamine (Caisson DML23) supplemented with 10% fetal bovine serum (Thermo Fisher 16000044). K562 cells were acquired from ATCC (CCL-243) and were maintained at 37°C with 5% CO_2_ in R10 media: RPMI with stabilized L-glutamine (Thermo Fisher 11875119) supplemented with 10% fetal bovine serum (Thermo Fisher 16000044). Cells were regularly passaged and tested for presence of mycoplasma contamination with MycoAlert Plus Mycoplasma Detection Kit (Lonza).

Lentivirus was produced by polyethylenimine linear MW 25000 (Polysciences 23966) transfection of HEK293FT cells with the transfer plasmid containing a Cas9 effector, or gRNA library, packaging plasmid psPAX2 (Addgene 12260) and envelope plasmid pMD2.G (Addgene 12259). After 72 h post-transfection, cell media containing lentiviral particles was harvested and filtered through 0.45 mm filter Steriflip-HV (Millipore SE1M003M00). K562 cells were transduced with KRAB-dCas9 or KRAB-dCas9-MeCP2 at a low multiplicity-of-infection (MOI < 1). Transduced K562 cells were selected with 10 ug/uL blasticidin (Thermo A1113903) for 10 days to enrich for expression of the dCas9 effector proteins. To isolate individual clones, K562 polyclonal lines were serially diluted to 50 cells per 10mL media. We then plated 100uL of this cell-media mix in 96-well round bottom plates (∼0.5 cells/well).

### Digital PCR for CRISPRi gene repression

We compared the CRISPRi v1 and v2 systems by targeting the transcription start sites and known enhancers of three genes (*MRPS23, SLC25A37* and *FSCN1)* with two gRNAs per targeted region. We synthesized gRNAs as top and bottom strand oligos (IDT) and cloned them into BsmBI-digested lentiGuideFE-Puro. We transduced the cells in biological triplicate with gRNA lentiviruses at a low MOI and after 24 hours selected for cells with gRNAs using puromycin (1 ug/uL, Thermo Fisher A1113803). We harvested the cells 10 days after transduction and extracted RNA using TRIzol (ThermoFisher 15596026). We quantified RNA concentration by spectrophotometry (NanoDrop). To measure gene expression, we performed digital PCR (Formulatrix Consellation) with Cy5/Iowa Black RQ target gene probes (IDT), FAM/ZEN/Iowa Black FQ for the actin normalizer (IDT), and Luna Universal One-Step RT qPCR Master Mix kit (NEB E3005L) and Tween-20 (Sigma-Aldrich P1379). We first normalized the target gene expression by actin expression per sample and then normalized this ratio to the ratio from cells transduced with non-targeting control gRNAs.

### CRISPR inhibition library design and cloning

We designed 20 nt gRNAs to target within 200 bp of the 88 selected plausibly causal noncoding variants. We used FlashFry v1.10.0 *(21)* to retain gRNAs with the lowest predicted off-target activity, as estimated by the Hsu-Scott score *(72)*. Each SNP was targeted by two different gRNAs. In addition, we also included in our library 12 non-targeting gRNAs from the GeCKOv2 library *(22)* as negative controls and 12 gRNAs targeting the TSSs of six non-essential genes as positive controls. The six non-essential genes (*CD46, CD52, HSPA8, NMU, PPIA* and *RPL22)* were identified by a CRISPR knock-out screen in K562 cells using the PICKLES database *(73)*. We additionally included 10 gRNAs targeting the CD55 TSS for our FACS-based multiplicity-of-infection (MOI) estimator, bringing the total number of gRNAs to 210.

To clone the gRNA library, top and bottom strand oligos (IDT) were resuspended in water at 100 uM and then mixed at 1:1 ratio for each gRNA. Then, 1uL of the oligo mix was added to a master mix containing 1x T4 ligase buffer (NEB M0202M), 0.5 uL T4 PNK (NEB M0201L) and water to a final concentration of 10uL per reaction. For oligo annealing, we incubated the oligo mix at 37°C for 30 min, then 95°C for 5 min with a temperature change of 1°C/5s until reaching 4°C. To create the oligo pool, we pooled together 3 uL of each annealed oligo. The oligo pool was diluted 1:10 with water and then cloned in the lentiGuideFE-Puro, which was linearized with BsmBI (Thermo ER0451) and dephosphorylated. The ligation was performed in 11 reactions with each reaction consisting of 5uL Rapid Ligation Buffer (Enzymatics B101), 0.5 uL T7 ligase (Enzymatics L602L), digested plasmid at 25 ng per reaction, 1 uL diluted oligo mix and ddH2O to final volume of 10 uL. The ligation was performed at room temperature for 15 min.

Next, 100 uL of the combined ligation reactions were mixed with 100 uL isopropanol, 1 uL GlycoBlue (Thermo Fisher AM9515) and 2 uL of 5M NaCl (50 mM final concentration), incubated for 15 min at room temperature, and spun at 12,000 g for 15 min. The pellet was washed twice with prechilled 70% ethanol, air dried for 15 min or until dried completely, resuspended in 5 uL 1x TE buffer (Sigma). Next, 2 uL of library ligation was added to 50 uL Endura cells (Lucigen) then electroporated, recovered and plated. The following day bacterial colonies were scraped, plasmids were isolated using a maxi prep (Qiagen 12965) and library representation was determined by MiSeq (Illumina).

To generate and concentrate the pooled library, lentivirus was generated as described above. Briefly, we seeded 10 × 225 cm^2^ flasks with HEK293FT cells and, at 70% confluency, we co-transfected the pooled gRNA library, psPAX2 and pMD2.G. Lentivirus was collected 72 hours post-transfection and filtered using a 0.45 um filter. The supernatant was then ultracentrifuged for 2 hr at 100,000 g (Sorvall Lynx 6000), and the pellet was resuspended overnight at 4°C in phosphate-buffered saline with 1% bovine serum albumin.

### Multiplicity-of-infection estimation via flow cytometry

When transducing cells at a high MOI, it is not possible to estimate the MOI by traditional methods (*e*.*g*. survival after drug selection) or without the time and cost of single-cell sequencing. By including multiple gRNAs that target the *CD55* TSS (10 gRNAs), we were able to estimate the number of gRNAs per cell (MOI) using flow cytometry for CD55 cell surface protein knockdown over a range of viral transduction volumes. We performed two transductions with concentrated lentivirus (4 uL and 6 uL) and, after 48 hours, we selected with puromycin for 10 days. We included three positive control transductions with different *CD55* TSS-targeting gRNAs and three negative control transductions with three different non-targeting gRNAs.

For flow cytometry, 1×10^6^ cells per condition were harvested and washed with PBS after selection. The cells were stained for 5 minutes at room temperature with LIVE/DEAD Fixable Violet Dead Stain Kit (ThermoFisher L34864). Subsequently, the cells were stained with antibodies for 20 minutes on ice with 1uL CD55-FITC (clone JS11) (BioLegend 311306). Cells were washed with PBS to remove unbound antibodies prior to sorting. Cell acquisition and sorting was performed using a Sony SH800S cell sorter. Sequential gating was performed as follows: 1) exclusion of debris based on forward and side scatter cell parameters, 2) dead cell exclusion. The sorting gates were set such that 90% of live K562 cells would be considered CD55 positive. We estimated that 6 uL viral volume yields an MOI of ∼13.5 and elected to use this condition for our STING-seq assay (**Fig. S2**).

### Expanded CRISPR-compatible Cellular Indexing of Transcriptomes and Epitopes (ECCITE-seq)

For ECCITE-seq, we ran one lane of a 10x Genomics 5’ kit (Chromium Single Cell Immune Profiling Solution v1.0, 1000014, 1000020, 1000151) targeting recovery of 20,000 cells per lane (superloading). We found this required loading approximately 44,000 cells to recover 21,300 total cells (including multiple cells per droplet counts, or “multiplets”). Cell hashing was performed as described in a previously published protocol using four hashtag-derived oligonucleotides (HTOs) using hyperconjugation *(24)*. Gene expression (cDNA), hashtags (HTOs) and guide RNA (Guide-derived oligos, GDOs) libraries were constructed by following 10x Genomics and ECCITE-seq protocols. We sequenced the cDNA, HTO and GDO libraries with two NextSeq 500 1×75 high-output runs (Illumina).

### Single cell data processing

After demultiplexing the sequencing reads, we generated UMI count matrices for the libraries using the kallisto indexing and tag extraction (kite) workflow with the hg38 reference genome (Ensembl v96), followed by the kallisto | bustools scRNA-seq pipeline *(74)*. We then analyzed the UMI count matrices in R v.4.0.2 with Seurat v.3.0.0 *(41)* and custom scripts for differential gene expression testing within the SCEPTRE framework *(26)*. We processed the cDNA UMI count matrix and retained cells with less than 20% mitochondrial reads and at least 850 unique gene UMIs. This resulted in 14,775 cell barcodes and 3,875 median genes per cell. We identified 14,728 cells in the HTO UMI count matrix that passed initial quality control of at least one HTO detected per cell and less than 2,500 total HTO UMIs per cell. We center-log-ratio (CLR) transformed the HTO UMI counts and demultiplexed cells by their transformed HTO counts to identify singlets. We used the HTODemux function and determined a threshold of 0.91 maximized the number of singlets detected to 9,520 while reducing the number of multiplets (multiple cells per droplet) to 2,482 and empty cells to 2,726. We used the 9,520 singlets to then process the gRNA (GDO) UMI count matrix, identifying 9,507 cells after removing cells with more than 15,000 total GDO UMIs. We observed that a minimum of 5 UMIs per gRNA resulted in a stable threshold as the number of cells per gRNA does not change with higher thresholds (**Fig. S3A**).

### Pairwise gRNA-to-gene tests with SCEPTRE

We generated raw UMI count matrices for gene expression and gRNA expression, along with accompanying single cell meta-data to use as covariates in model fitting (**Table S3B**). We defined for each gRNA a list of genes within 500 kb to be tested for differential expression in *cis*. For each gRNA-to-gene pair, we extracted that gene’s UMI counts and labeled the cells with a given gRNA. We then tested for differential expression within the SCEPTRE framework *(26)*, adjusting for the following single cell covariates: total gene expression UMIs, unique genes, total gRNA expression UMIs, unique gRNAs and percentage of mitochondrial genes. Briefly, SCEPTRE is a statistical framework to analyze high MOI CRISPR screens in single cells with state-of-the-art calibration. First, SCEPTRE fits a negative binomial distribution to measure the effect of a single gRNA on a given gene via *Z*-score. Then, the distribution of gRNAs to cells is randomly sampled to build a gRNA-specific null distribution, recomputing a negative binomial *Z*-score. A skew-*t* distribution is fit to compare the test *Z*-score and the null distribution, and a two-sided *p*-value is derived, allowing for significance tests of increased or decreased gene expression *(26)*. To test for differential expression in *trans*, we defined for each gRNA a list of all 10,325 genes detected and repeated the test above. Non-targeting gRNAs were tested against all genes used in the *cis* and *trans* settings discussed previously and randomly sampled to match the number of *cis* and *trans* targeting gRNA-to-gene tests displayed on QQ-plots.

### Independent replication of trans-effect genes

Genomic regions corresponding to the CREs identified by rs524137 (*GFI1B*) and rs79755767 (*NFE2)* were targeted in two separate studies using the CRISPRi v1 system (dCas9-KRAB) in K562 cells, featuring different gRNAs and 10x Genomics 3’ gene expression *(7, 8)*, as opposed to the 5’ gene expression workflow we implemented. We re-analyzed both studies for replication of our identified target genes in *trans*, and defined replication as a SCEPTRE *p*-value < 0.05 and a direction of effect consistent with what we observed in our study. Re-analyzing data from Gasperini *et al*. *(7)*, we tested 569 genes identified from perturbing the rs524137-*GFI1B* CRE and replicated 321 genes (including *GFI1B*). Re-analyzing data from Xie *et al*. *(8)*, we tested 134 genes identified from perturbing the rs79755767-NFE2 CRE and replicated 47 genes (including *NFE2)*.

### Trans-regulated network gene set enrichments

We used two previously published chromatin immunoprecipitation sequencing (ChIP-seq) datasets in K562 cells to identify GFI1B *(35)* and NFE2 *(38)* transcription factor binding sites. To find genes with proximal ChIP-seq peaks, we defined a 1 kb window around all protein-coding gene TSSs and used bedtools v2.25.0 *(67)*. To test for enrichment of ChIP-seq peaks in *GFI1B* or *NFE2 trans*-regulatory gene sets, we constructed contingency tables and used Fisher’s exact test to test for significance and compute odds ratios with 95% confidence intervals. To construct GWAS-identified sets of genes, we used all fine-mapped SNPs from the 29 GWASs previously described (categorized by cell type) with a high Bayes factor for being plausibly causal (log_10_ BF ≥ 2). GWAS gene enrichment was performed in a similar fashion as for ChIP-seq peaks.

### Gene co-expression analyses and bone marrow single cell gene expression

To compute co-expression matrices for each *trans*-regulatory network, we used cDNA UMI count matrices with missing genes per cell imputed with MAGIC *(75)*. As a measure of co-expression, the biweight midcorrelation, a weighted correlation analysis, was calculated for each pair of genes *(76)*. Genes were then clustered based on their co-expression patterns by hierarchical clustering. Transcription factor binding site and GWAS gene enrichment was performed as described above. We used Human Cell Atlas single cell RNA-sequencing from 35 bone marrow donors *(40)* and identified 27 cell types as described previously *(41)*. Single cell data were processed with Seurat v.4.0.0 *(77)* to generate UMAP plots and heatmaps. To visualize entire *trans*-regulatory network clusters on a UMAP plot, we plotted the mean expression of all cluster genes within each cell.

### STING-seq power calculations for other GWASs

We down-sampled the *trans*-regulatory network gRNA-to-gene tests for both rs524137 targeting gRNAs and the top rs73660574 and rs79755767 targeting gRNAs. All of these gRNAs were recovered in sufficient (high) numbers of single cells (923, 877, 841 and 761, respectively). For each gRNA-to-gene pair, we randomly sampled the number of cells tested with each gRNA from 90% to 10% by 10% increments, repeating each step 100 times. We then compared the mean down-sampled SCEPTRE *p*-value for each step with the nominal significance threshold (1% FDR) for each gRNA’s *trans*-effects tests to classify whether it retained significance. We then divided all gRNA-to-gene pairs by their expression level and log_2_ fold-change into tertiles to examine at what number of cells could 80% of significant findings be retained.

To perform GWAS projections, we identified the number of independent GWAS loci from five studies: BMI-adjusted waist-hip ratio *(42)*, type 2 diabetes *(43)*, schizophrenia *(44)*, rheumatoid arthritis *(45)* and stroke *(46)*. We assumed that statistical fine-mapping would be successful for each of these loci and that a single noncoding variant would be targeted, allowing us to estimate the number of single cells needed for STING-seq. We allowed for two gRNAs to target each variant and for 10 gRNAs per cell, then computed the number of single cells required for 80% power based on the down-sampled analyses for moderately-expressed (middle tertile) genes.

