## Supplementary Figures for "Discovery of target genes and pathways of blood trait loci using pooled CRISPR screens and single cell RNA sequencing"

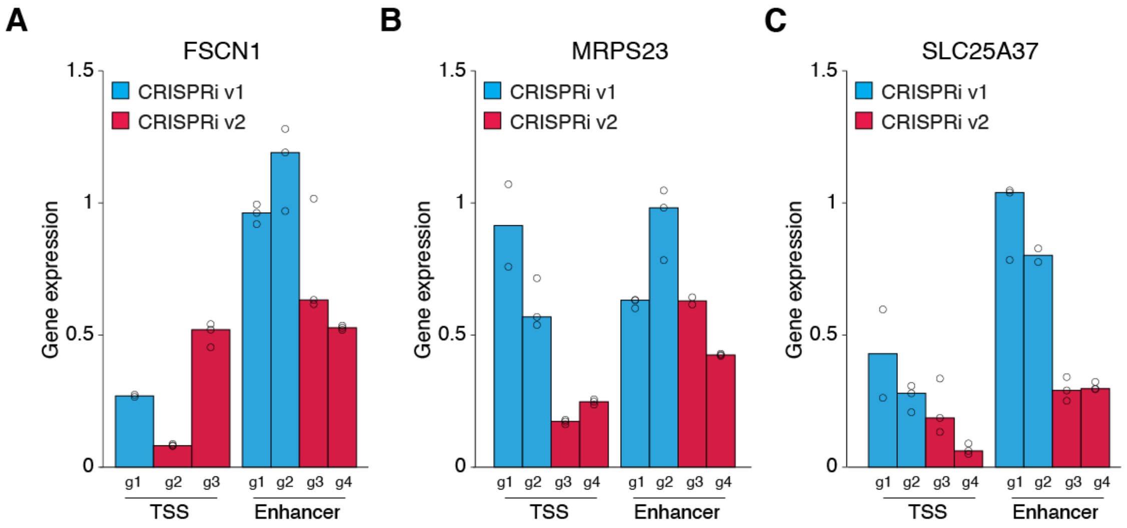


**Figure S1. Digital PCR for CRISPR inhibition (CRISPRi) with individual genes and guide RNAs.**

Digital PCR gene expression in K562 by targeting the transcription start sites (TSS) and known enhancers of *FSCN1* (**A**), *MRPS23* (**B**) and *SLC25A37* (**C**) with either CRISPRi v1 or v2. Each bar represents one gRNA ($n$ = 3 biological replicates per gRNA).

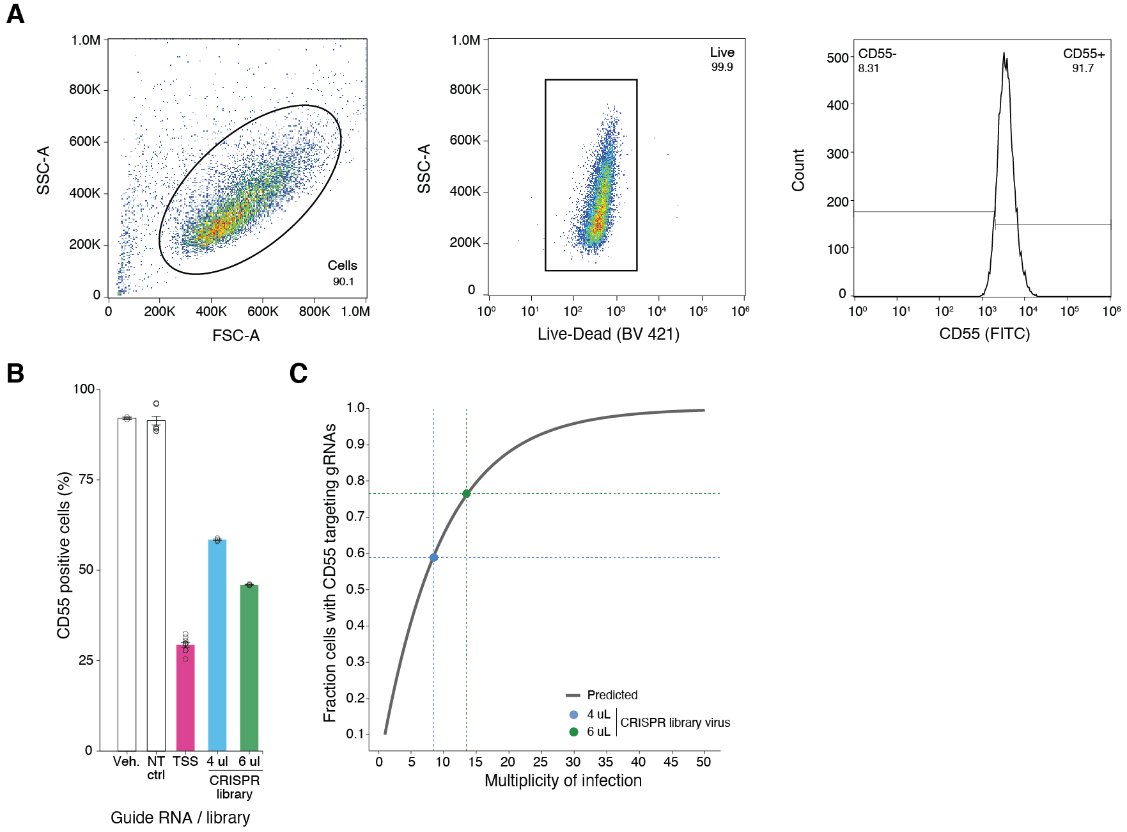


**Figure S2. Flow cytometry for estimation of multiplicity of infection.**

(**A**) Flow cytometry gating strategy to quantify cell surface expression of CD55. Live cells were gated by the forward and side scatter area then viable cells were selected by gating on side scatter area and LIVE/DEAD Violet. Sorting gates were set so that 90% of wild-type K562 cells without any guide RNA (gRNA) are classified as CD55 positive. (**B**) CD55 positive cells with no transduction (veh.), non-targeting gRNAs, transcription start site (TSS)-targeting gRNAs and two separate volumes of the STING-seq pooled library. (**C**) Estimation of the multiplicity-of-infection based on the starting distribution of *CD55* TSS-targeting gRNAs and proportion of cells bearing a gRNA.

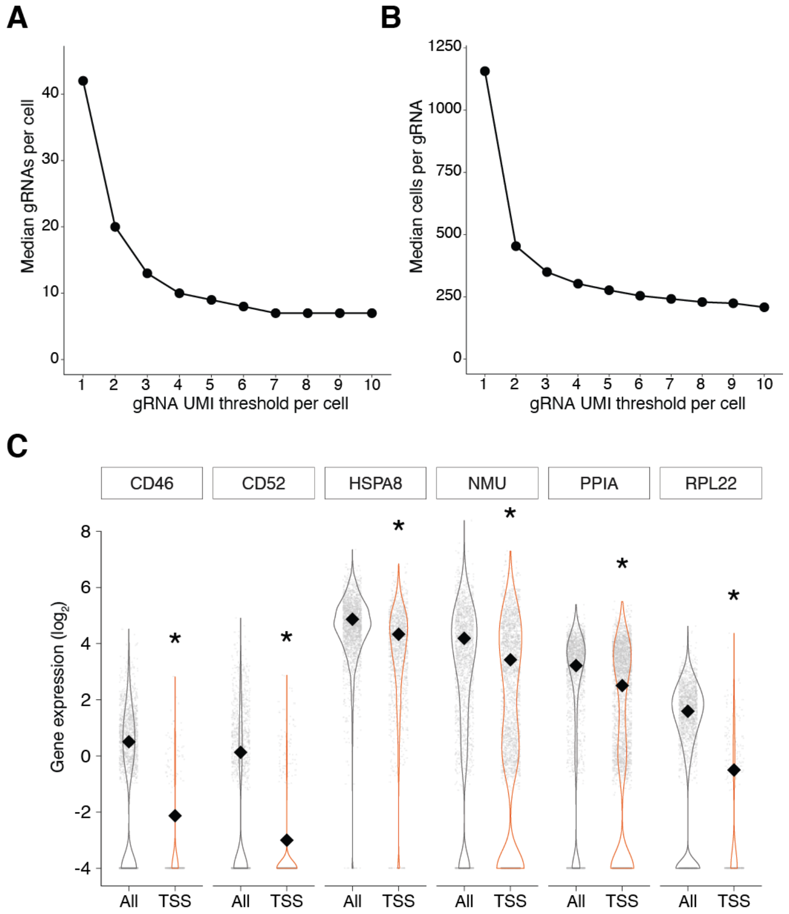


**Figure S3. Unique molecular identifier (UMI) thresholds for guide RNA (gRNA) detection and gene expression for positive controls.**

(**A**) Median number of gRNAs per cell detected at different minimum gRNA UMI thresholds. (**B**) Median number of cells per gRNA detected at different minimum gRNA UMI thresholds. A gRNA UMI threshold of 5 was selected for assigning gRNAs to cells. (**C**) Normalized single cell gene expression for the top transcription start site-targeting gRNAs for six positive controls, *CD46*, *CD52*, *HSPA8*, *NMU*, *PPIA* and *RPL22*. * indicates $p < 0.05$ (SCEPTRE $p$-value with Bonferroni correction).

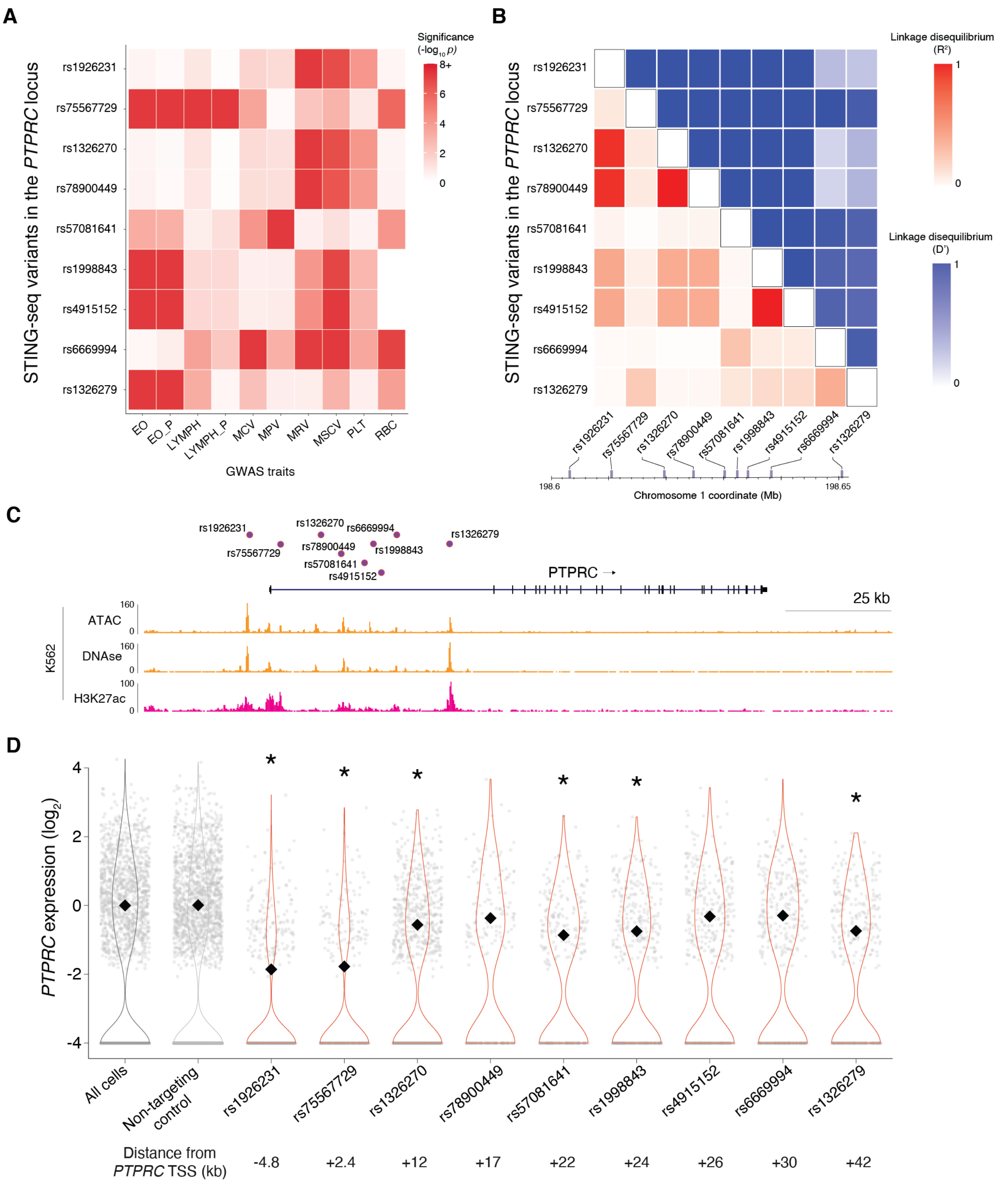


**Figure S4. STING-seq of nine GWAS variants at the *PTPRC* locus.**
(**A**) Heatmap of *p*-values from 10 blood trait GWASs for nine variants mapping to cCREs proximal to *PTPRC*. The maximal color value indicates genome-wide significance ($5x10^{-8}$). (**B**). Pairwise linkage disequilibrium matrix ($R^2$ and D') for the nine targeted variants using the 1000 Genomes

CEU and GBR populations. (**C**) Normalized single cell *PTPRC* expression for the top variant-targeting gRNAs. *PTPRC* was differentially expressed upon perturbation of six out of nine variants, identifying six significant CREs (5% FDR). Two variants (rs1926231 and rs75567729) are located closest to the TSS of the *PTPRC* gene and have the strongest impact on gene expression. However, they are not in LD and have different GWAS significance patterns. Two other variants (rs78900449 and rs4915152) were not significant but were in strong LD ($R^2 \geq 0.95$) with significant variant-identified CREs for *PTPRC* (rs1326270 and rs1998843, respectively), suggesting they may be non-functional LD proxy variants.

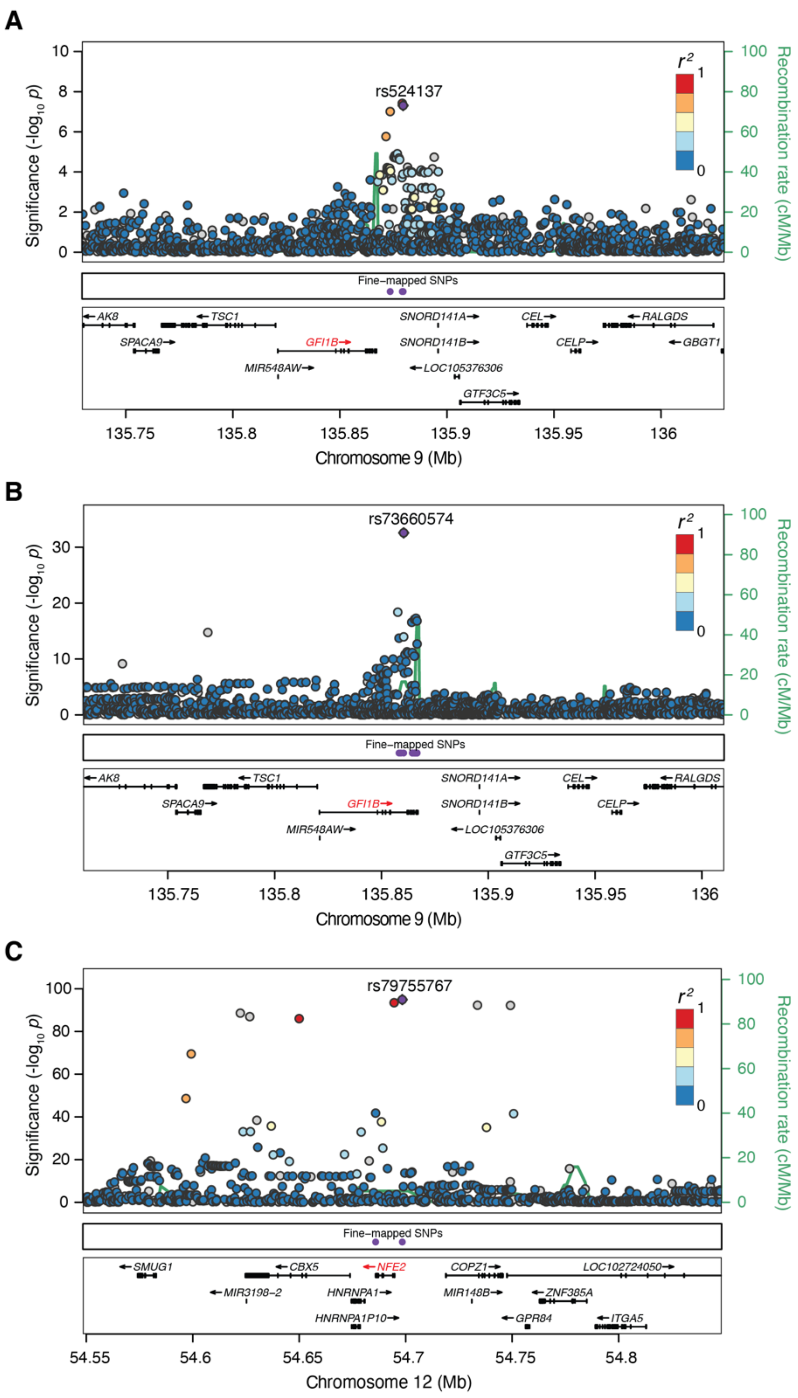


**Figure S5. GWAS association of STING-seq variants with significant *trans* gene expression changes.**

(**A**) LocusZoom plot of a monocyte percentage locus where rs524137 was targeted. (**B**) LocusZoom plot of a mean reticulocyte volume locus where rs73660574 was targeted. (**C**) LocusZoom plot of a red blood cell distribution width locus where rs79755767 was targeted.

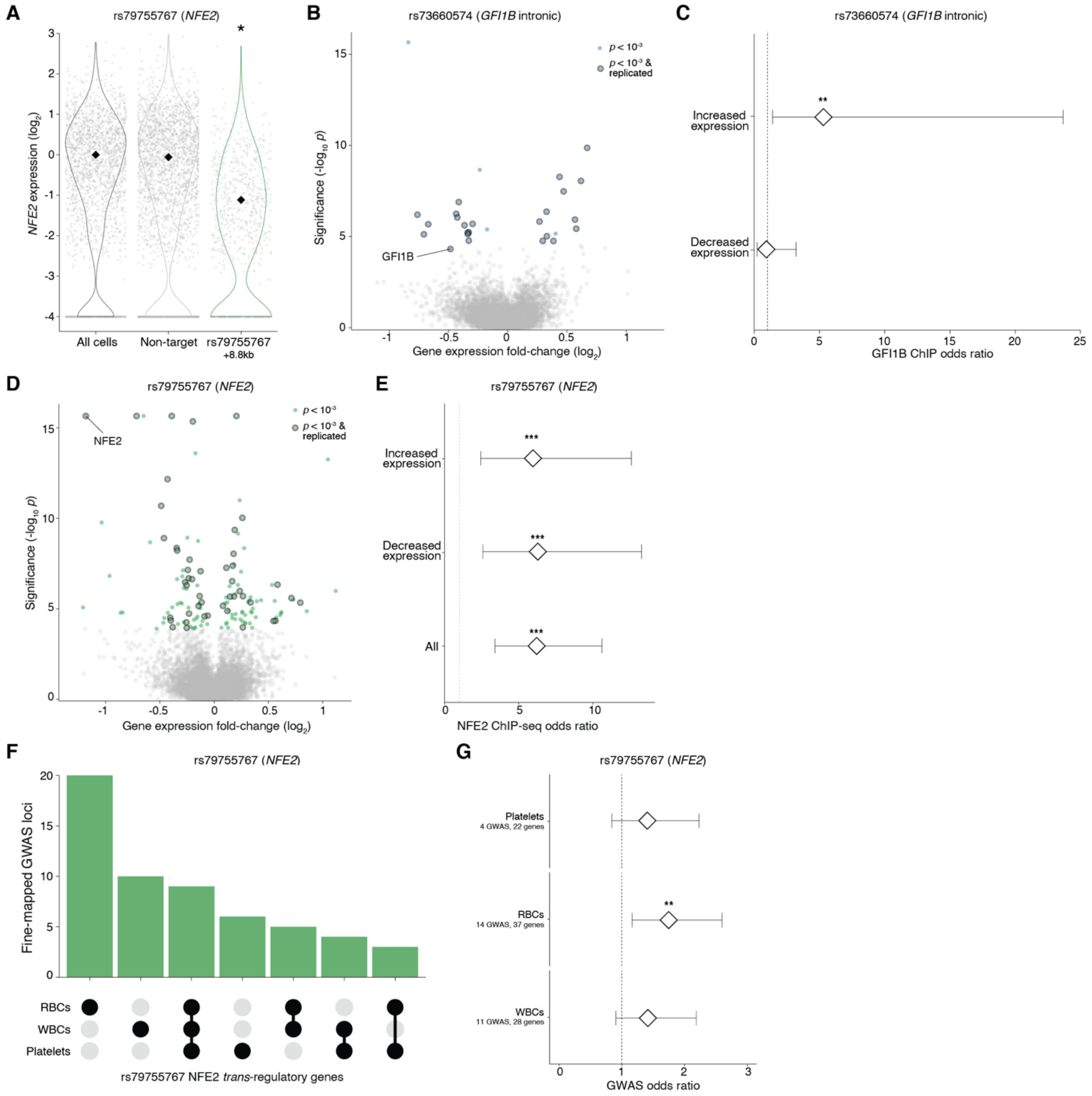


**Figure S6. Analysis of *trans*-regulated genes for rs79755767 and rs73660574.**
(**A**) Normalized gene expression of *NFE2* in all cells, cells receiving a non-targeting guide RNA (gRNA) and cells receiving a gRNA that targets the *cis*-regulatory element (CRE) that overlaps rs79755767. (**B**) Volcano plot of top rs73660574-*GFI1B trans*-regulated gRNA-to-gene pairs. rs73660574 is the weaker, intronic *GFI1B* enhancer. 28 unique genes were identified for the rs73660574-*GFI1B trans*-regulatory network, 24 of which were identified in the rs524137-*GFI1B* network. (**C**) rs73660574-*GFI1B trans*-regulated genes with increased expression were enriched for GFI1B ChIP-seq peaks (OR = 5.22, $p = 5.6\text{x}10^{-3}$). (**D**) Volcano plot of top rs79755767-*NFE2*

*trans*-regulated gRNA-to-gene pairs in *trans*. 134 unique genes were identified for the rs79755767-*NFE2 trans*-regulatory network. (**E**) rs79755767-*NFE2 trans*-regulatory genes with increased expression were enriched for NFE2 ChIP-seq peaks. (**F**) rs79755767-*NFE2 trans*-regulated genes overlapped genes closest to fine-mapped GWAS variants, grouped by the cell type of the GWAS trait (platelets, white blood cells or red blood cells). Only genes with an overlap are shown. (**G**) rs79755767-*NFE2 trans*-regulated genes were enriched for red blood cell genes (OR = 1.76, $p$ = 6.2x$10^{-3}$).

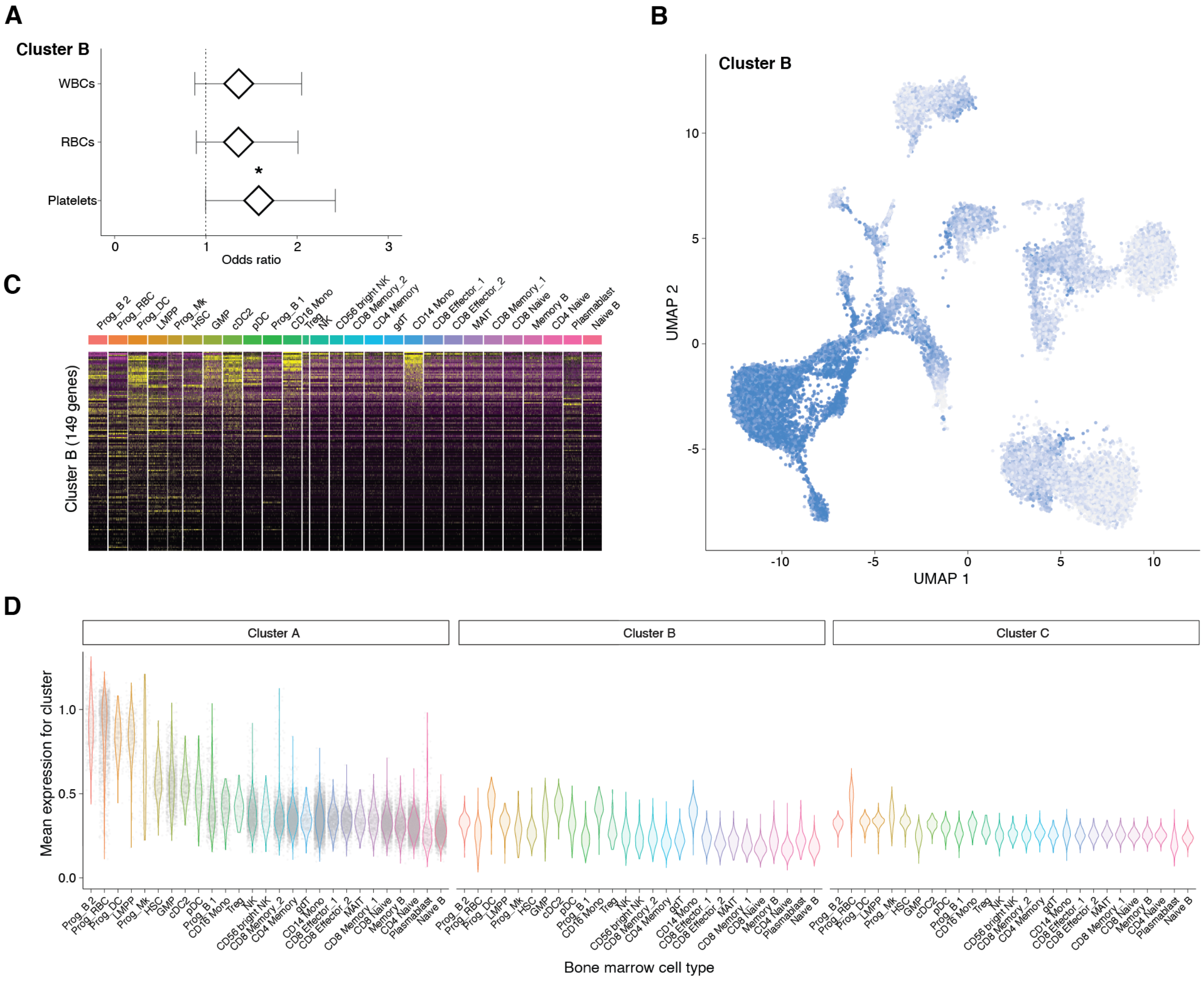


**Figure S7. rs524137-*GFI1B trans*-regulated cluster B and mean expression in human bone marrow for all clusters by cell type.**

(**A**) rs524137-*GFI1B trans*-regulatory network cluster B enrichment for genes mapping closest to fine-mapped variants for WBC, RBC and platelet GWAS traits. (**B**) Mean expression of cluster B genes per single cell, projected onto the bone marrow UMAP plot. The cell types with the highest Cluster B expression patterns are differentiated cells, whereas expression in bone marrow progenitor cells is lower. (**C**) Expression of genes in cluster B ($n$ = 151 genes) by cell type in 35 bone marrow donors from the Human Cell Atlas. (**D**) Mean expression of genes included in each cluster (clusters A, B and C) in each bone marrow cell type. * indicates $p < 0.05$.

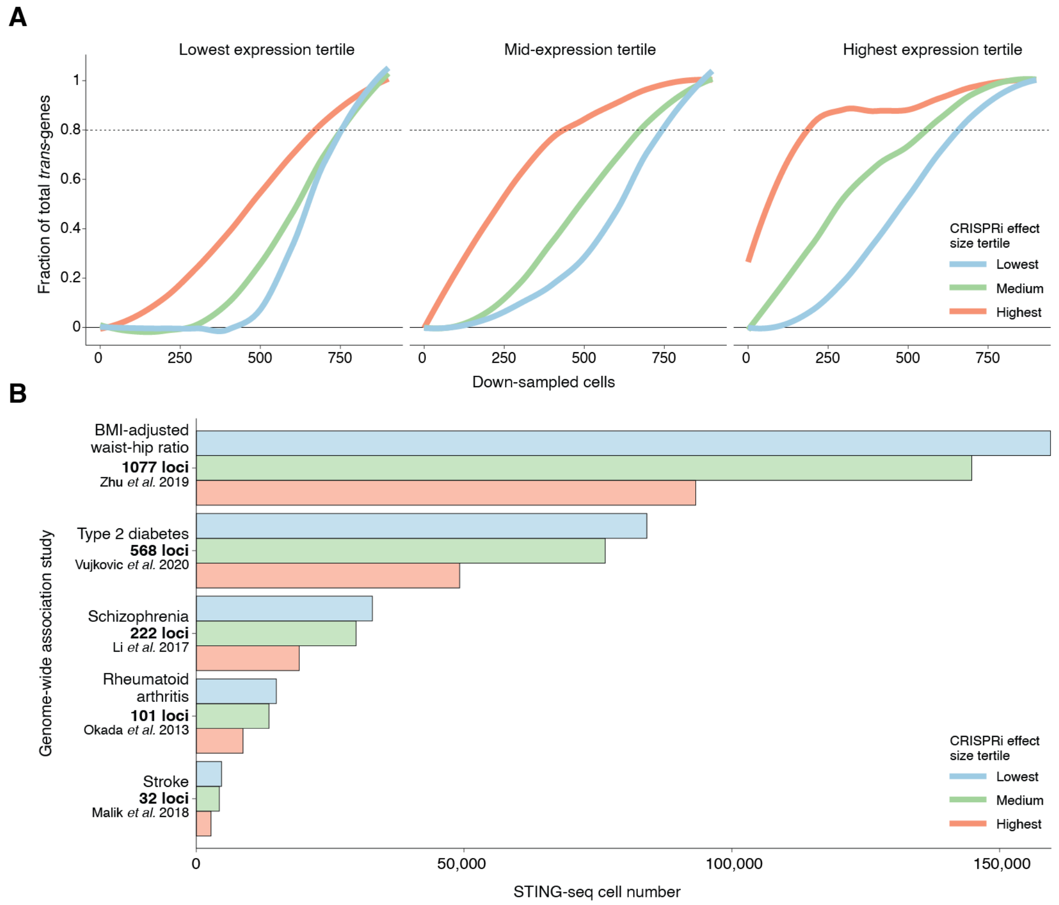


**Figure S8. Power estimation for application of STING-seq to other traits and diseases.** (**A**) Power analysis of single cells needed to detect changes in gene expression in *trans*. *Trans* effects are divided into tertiles based on gene expression and, separately, CRISPR perturbation-induced changes in expression are divided into tertiles. (**B**) Single cells needed to perform STING-seq (80% power) at one variant per locus for BMI-adjusted waist-hip ratio (1077 loci), type 2 diabetes (568 loci), schizophrenia (222 loci), rheumatoid arthritis (101 loci) and stroke (32 loci).
